# Aging Differentially Alters the Transcriptome and Landscape of Chromatin Accessibility in the Male and Female Mouse Hippocampus

**DOI:** 10.1101/2023.10.17.562606

**Authors:** Jennifer M. Achiro, Yang Tao, Fuying Gao, Chia-Ho Lin, Marika Watanabe, Sylvia Neumann, Giovanni Coppola, Douglas L. Black, Kelsey C. Martin

**Affiliations:** Department of Biological Chemistry, David Geffen School of Medicine, UCLA; Department of Psychiatry and Biobehavioral Sciences and Semel Institute for Neuroscience and Human Behavior, David Geffen School of Medicine, UCLA; Department of Microbiology, Immunology and Molecular Genetics, UCLA

**Keywords:** Aging, sex differences, sex-bias, hippocampus, gene expression, alternative splicing, synaptic proteins, myelin sheath, ATAC-seq, chromatin accessibility, LINE-1, retrotransposable elements derived sequences, male-biased promoter chromatin accessibility

## Abstract

Aging-related memory impairment and pathological memory disorders such as Alzheimer’s disease differ between males and females, and yet little is known about how aging-related changes in the transcriptome and chromatin environment differ between sexes in the hippocampus. To investigate this question, we compared the chromatin accessibility landscape and gene expression/alternative splicing pattern of young adult and aged mouse hippocampus in both males and females using ATAC-seq and RNA-seq. We detected significant aging-dependent changes in the expression of genes involved in immune response and synaptic function, and aging-dependent changes in the alternative splicing of myelin sheath genes. We found significant sex-bias in the expression and alternative splicing of hundreds of genes, including aging-dependent female-biased expression of myelin sheath genes and aging-dependent male-biased expression of genes involved in synaptic function. Aging was associated with increased chromatin accessibility in both male and female hippocampus, especially in repetitive elements, and with an increase in LINE-1 transcription. We detected significant sex-bias in chromatin accessibility in both autosomes and the X chromosome, with male-biased accessibility enriched at promoters and CpG-rich regions. Sex differences in gene expression and chromatin accessibility were amplified with aging, findings that may shed light on sex differences in aging-related and pathological memory loss.

**Highlights:**

- Aging amplifies sex differences in the transcriptome and chromatin accessibility of mouse hippocampus
- Aging is associated with widespread changes in the transcriptome of mouse hippocampus, including in genes involved in immune response and synaptic function
- Myelin sheath-related genes in aged hippocampus show female-biased expression
- Alternative splicing of myelin sheath-related genes changes with aging in the hippocampus
- Sex-bias is evident in alternative splicing of nuclear and primary cilia genes
- Aging in the hippocampus is associated with increased chromatin accessibility in both males and females
- LINE-1 derived sequences are more accessible and LINE-1 transcripts are upregulated in the aged hippocampus of both males and females
- Promoter and CpG-rich regions show male-biased accessibility

## Introduction

Aging is associated with cognitive decline and memory impairment, including spatial and episodic memory deficits^1^. Research has revealed sex differences in memory performance with aging, as well as sex differences in the incidence of Alzheimer’s disease, a condition associated with memory loss and damage to the hippocampal brain region^2–6^. The hippocampus is critical for spatial, episodic and long-term memory formation^7^, and therefore understanding how sex differences impact aging in the hippocampus can provide insights into aging- and sex-dependent declines in memory.

Numerous studies have shown that the abundance of cell types in the brain remains largely constant during aging^8–12^, however increased immune activation, blood brain barrier breakdown, altered synaptic function, as well as changes in gene expression, alternative splicing, epigenetic marks and chromatin accessibility have been associated with aging in the brain^8,11,13–18^. Regulated chromatin accessibility is critical to the precise control of gene expression patterns^19^, and hippocampus-dependent long-term memory formation requires new gene expression^20^. However, few studies have examined sex-differences in aging-related changes in gene expression in the hippocampus^21,22^, and none, to our knowledge, have examined how sex differences affect aging-related alternative splicing and chromatin accessibility in the hippocampus. Understanding how gene regulatory networks and chromatin dynamics associated with aging differ between sexes in the hippocampus may guide sex-dependent therapies for age-related memory disorders.

## Materials and Methods

### Animals

Male and female C57B16/J mice, aged 8 weeks and 78 weeks, were purchased from Jackson Laboratories. Use of mice in this study was approved by the UCLA Institutional Animal Care and Use Committee. Mice were housed in groups under a 12:12 hr light/dark cycle with food provided ad libitum until they reached the age of 10 weeks (young adult) and 80 weeks (aged). For sample collection, mice were deeply anesthetized with isoflurane and euthanized by cervical dislocation. The brain was quickly removed and cooled for one minute in ice-cold PBS before the hippocampus from each hemisphere was dissected on ice and processed immediately. For sequencing experiments, one hippocampus was homogenized in Trizol for RNA preparation and the other hippocampus was homogenized for nuclei preparation for ATAC-seq (see below). For qPCR and western blot validation, hippocampal tissue was homogenized in Trizol or RIPA buffer, respectively (see below).

### RNA-seq Library Preparation and Sequencing

RNA was extracted from Trizol using a Trizol/RNeasy hybrid protocol. Briefly, after phase separation, the aqueous phase was mixed with one volume of 70% ethanol and passed through an RNeasy spin column (QIAGEN RNeasy Mini Kit). RNA quality was determined using the High Sensitivity RNA ScreenTape Assay (Agilent 5067-5579), and all samples had RIN scores > 7.0. RNA-seq libraries for differential expression or splicing analysis were prepared for four biological replicates, including four young adult males, four aged males, four young adult females, and four aged females. For differential expression analysis, total RNA libraries were prepared using the TruSeq Stranded Total RNA Library Prep Kit with Ribo-Zero (Illumina 20020596). Paired-end 100 bp sequencing to a depth of ∼50 million reads per sample was performed using the Illumina HiSeq 2500 system at the UCLA Broad Stem Cell Research Center Sequencing Core. For splicing analysis, mRNA libraries were prepared with poly-A selection using the TruSeq Stranded mRNA Library Prep Kit (Illumina 20020594) and paired-end 75 bp sequencing to a depth of ∼50 million reads per sample was performed using the Illumina HiSeq 2500 system at the UCLA Neuroscience Genomics Core.

### ATAC-seq Library Preparation and Sequencing

ATAC-seq sample preparation was performed for eight replicates according to detailed protocols obtained from Hongjun Song’s lab at the University of Pennsylvania^19,23^. Briefly, one hippocampus was homogenized with a Dounce tissue grinder (Wheaton 357544) in 2 mL HB buffer (1 mM DTT, 0.15 mM spermine, 0.5 mM spermidine, protease inhibitor (Sigma-Aldrich 04693159001), 0.3% IGEPAL-630, 0.25 M sucrose, 25 mM MgCl_2_, 20 mM Tricine-KOH). The homogenate was then filtered through a 40 μm strainer. The filtrate was centrifuged over one volume of cushion buffer (0.5 mM MgCl_2_, 0.5 mM DTT, protease inhibitor, 0.88 M sucrose) at 2800 g for 10 minutes in a swinging bucket centrifuge at 4°C. The nuclei pellet was resuspended in 20 μL PBS. 1 μL of the nuclei sample was stained with Hoechst 33342 to calculate nuclei concentration, and 50,000 nuclei per sample were used for downstream library preparation. Libraries were prepared using the Nextera DNA Library Prep kit (Illumina FC-121-1030) and quality was analyzed using the D1000 ScreenTape Assay (Agilent 5067-5582). Paired-end 50 bp sequencing to a depth of ∼40 million reads per sample was performed using the Illumina HiSeq 2500 system at the UCLA Broad Stem Cell Research Center Sequencing Core.

### RNA-seq Data Analysis

RNA-seq reads were mapped to the mouse genome (mm10) using STAR with the two-pass option^24^. Only uniquely mapped reads were used for downstream analysis. Lowly expressed genes were removed by retaining only genes with counts per million > 0.1 in at least four samples (17,821 genes). Read counts were normalized via the trimmed mean method before differential expression analysis (DEA)^25^. For DEA, the analysis package edgeR^26^ was used, with an FDR < 0.05 cutoff. For age-dependent expression differences, data from female and male samples were analyzed separately. For differential splicing analysis, the analysis package rMATS version 4.0.2^27^ was used. Alternative splicing events were included for analysis if events had FDR-adjusted p-value < 0.05, total reads ≥ 50 and skipping/junction reads > 20. For gene ontology enrichment analysis, the online tool DAVID (Database for Annotation, Visualization and Integrated Discovery) was used^28,29^. RNA-sequencing data have been deposited in NCBI’s Gene Expression Omnibus^30^ and are accessible through GEO Series accession number GSE244506 (https://www.ncbi.nlm.nih.gov/geo/query/acc.cgi?acc=GSE244506).

### ATAC-seq Data Analysis

Eight replicates per condition were sequenced for ATAC-seq; however, one young male sample had less than one million reads and so was not used in further analyses. Adapter content was trimmed using Scythe (https://github.com/vsbuffalo/scythe/) and ATAC-seq reads were mapped to the mouse genome (mm10) using Bowtie 2^31^. Reads in blacklist regions^32,33^ and reads mapped to mitochondria were removed. Peaks were called on each individual sample using MACS3 (with parameter settings --nomodel -f BAMPE and FDR cutoff of 0.05)^34^. For each sample, read pairs (fragments) were counted for each peak using featureCount^35^ (with parameter settings -p --countReadPairs -M --fraction). For quality control, we calculated a transcription start site (TSS) enrichment score using ATACseqQC^36^ and a FRiP score (fraction of reads/fragments in peaks). One young male and one aged male sample had a TSS enrichment score < 10 and/or a FRiP score < 0.2, therefore these samples were excluded from further analyses. The remaining samples had an average TSS enrichment score of 20.2 ± 1.1 (mean ± s.e.m.) and an average FRiP score of 0.46 ± 0.03 (mean ± s.e.m.). For each condition, narrow peak sets were merged, and peaks were included in consensus peak sets for each condition if there was at least a 50% overlap with a peak in at least two replicates (consensus sets: 65,088 peaks in young male, 71,719 in aged male, 61,264 in young female and 68,655 in aged female). A total consensus peak set (92,233 peaks) was generated by merging the consensus peak sets from each condition. Fragments were counted for each total consensus peak in each sample using featureCount as described above, and differential analysis was performed using DESeq2, including sequencing batch as a variable^37^. Batch correction for principal components analysis was performed using limma^38^. Peak annotation, peak histograms and motif searches were done using HOMER^39^. Browser track examples were generated from the UCSC genome browser (http://genome.ucsc.edu)^40^. For the ATAC-seq peak profile heat map, normalized peak histograms for each peak were generated in HOMER for each sample using 10 bp bins. Then for each peak, an average ATAC histogram was calculated for each condition and ranked by young male ATAC signal intensity +/- 100 bp around peak center. For ATAC-seq profile plots, normalized peak histograms for each sample were generated in HOMER using 10 bp bins, producing one histogram per sample based on the number of peaks (or gene TSSs) indicated in figures, and then averages and standard error of the means were calculated based on the sample histograms (with n = number of samples, i.e. 6 young male, 7 aged male, 8 young female, 8 aged female). Genomic locations of full-length intact long interspersed element-1 (LINE-1) regions were obtained using L1Base 2^41^. Where indicated in the results, we used smooth-quantile with GC-content normalization (QSmooth-GC) using qsmooth^42,43^. This work used computational and storage services associated with the Hoffman2 Shared Cluster provided by UCLA Institute for Digital Research and Education’s Research Technology Group. ATAC-sequencing data have been deposited in NCBI’s Gene Expression Omnibus^30^ and are accessible through GEO Series accession number GSE244506 (https://www.ncbi.nlm.nih.gov/geo/query/acc.cgi?acc=GSE244506).

### qPCR

Samples for qPCR validation were prepared from separate animals than those used for library preparation. Fifty ng of total RNA was reverse transcribed into cDNA using SuperScript III First Strand Synthesis System (Invitrogen 18080051) with random hexamer primers for four biological replicates of each sex and age group. Technical triplicates were prepared for each primer set using SYBR Green PCR Master Mix (Applied Biosystems 4309155). The following primers were used:

\**Gapdh* (F: AGGTCGGTGTGAACGGATTTG; R: TGTAGACCATGTAGTTGAGGTCA),

\**Tubb3* (F: TAGACCCCAGCGGCAACTAT; R: GTTCCAGGTTCCAAGTCCACC),

\**Gfap* (F: CCCTGGCTCGTGTGGATTT; R: GACCGATACCACTCCTCTGTC),

\**Pcdhb9* (F: ACTGCTCTTGAGAATACCAGAGA; R: AGGACGTGAAAATAAGGGTTGG),

*Gpr17* (F: TCACAGCTTACCTGCTTCCC; R: CCGTTCATCTTGTGGCTCTTG),

\**Ptpro* (F: AACATCCTGCCGTATGACTTTAG; R: GGGACTTCTGTTGTAGGACCATC),

\**Npnt* (F: GGACAGGTCCGATGTCAGTG; R: CTTCCAGTCGCACATTCATCA),

\**Mag* (F: CTGCCGCTGTTTTGGATAATGA; R: CATCGGGGAAGTCGAAACGG),

\**Mbp* (F: GACCATCCAAGAAGACCCCAC; R: GCCATAATGGGTAGTTCTCGTGT), LINE-1 5’UTR (F: TGAGTGGAACACAACTTCTGC; R: CAGGCAAGCTCTCTTCTTGC), LINE-1 ORF1 (F: ATGGCGAAAGGCAAACGTAAG; R: ATTTTCGGTTGTGTTGGGGTG),

LINE-1 5’UTR:ORF1 (F: CTGCCTTGCAAGAAGAGAGC; R: AGTGCTGCGTTCTGATGATG).

Gene primers from PrimerBank (http://pga.mgh.harvard.edu/primerbank/) are indicated with a *. Primers for LINE-1 were obtained from previous publications^44,45^. The rest of the gene primers were custom-designed to span exon-exon junctions. The samples were run on the Bio-Rad CFX Connect Real-Time PCR Detection System. We used the delta-delta Ct method for calculating fold gene expression, with delta Cts calculated relative to *Gapdh* (for RNA-seq validation) or *Tubb3* (for LINE-1), and delta delta Cts calculated relative to young male samples. Results were tested for normality using the Shapiro-Wilk test. Unpaired t-tests/one-way ANOVAs with Sidak’s multiple comparisons test or Mann-Whitney/Kruskal-Wallis tests with Dunn’s multiple comparison tests were used for normally and non-normally distributed data, respectively.

### Western Blot

Western blot samples were prepared from four replicates. For each replicate, one hippocampus was homogenized in RIPA buffer with protease and phosphatase inhibitors and centrifuged at 10,000 g for 10 min at 4°C to remove cell debris. Protein concentration was determined using the Pierce BCA Protein Assay Kit (Thermo Fisher 23225). All protein samples were diluted to a protein concentration of 0.5 mg/mL. 4X sample loading buffer (0.2 M Tris-HCl pH 6.5, 4.3 M glycerol, 8.0% (w/v) SDS, 6 mM bromophenol blue, 0.4 M DTT) was added and the samples were boiled at 95-100°C for 10 min. The samples were then aliquoted and stored at -20°C. Samples were loaded on NuPAGE 4-12% Bis-Tris protein gels (Invitrogen NP0335BOX) and transferred onto 0.2 μm nitrocellulose membranes. After transfer, blots were blocked with Odyssey TBS blocking buffer (LI-COR 927-50000) for 1 hr at room temperature before being incubated in primary antibody at 4°C overnight. The following primary antibodies were used: anti-MBP (Abcam, ab218011) and anti-TUJ1 (Biolegend, 801201). Blots were incubated in secondary antibodies for 2 hr at room temperature. Blots were imaged using the LI-COR Odyssey imaging system and quantified using LI-COR Image Studio software.

## Results

To investigate sex differences in aging-related changes in the transcriptome (including alternative splicing) and chromatin accessibility in the hippocampus, we dissected the hippocampus from each hemisphere of young adult (10 weeks) or aged (80 weeks) mice of both sexes and used one hippocampus for RNA-sequencing (RNA-seq) and one hippocampus for Assay for Transposases-Accessible Chromatin sequencing (ATAC-seq)^23,46^ library preparation (Fig. 1A). Principle component analysis of the RNA-seq data showed that samples were well separated by sex and age by the first and second principal component, respectively (Fig. 1B). Overall, 905 genes were differentially expressed (DE) either by sex or aging, with the largest number of differences occurring with aging irrespective of sex (Fig. 1C; Supp. Table 1). In the hippocampus of females, we found 446 genes that were upregulated and 261 genes downregulated with aging. Significantly fewer aging-associated changes were detected in the hippocampus of males compared to females (chi-square test, p<0.001, *X*^2^ = 108.1), with 206 and 164 genes being upregulated and downregulated in males, respectively, with aging (Fig. 1D, E). These results indicate that aging is associated with widespread changes in gene expression, with a larger number of genes undergoing aging-related changes in expression in the hippocampus of females than males.

**Figure 1:**
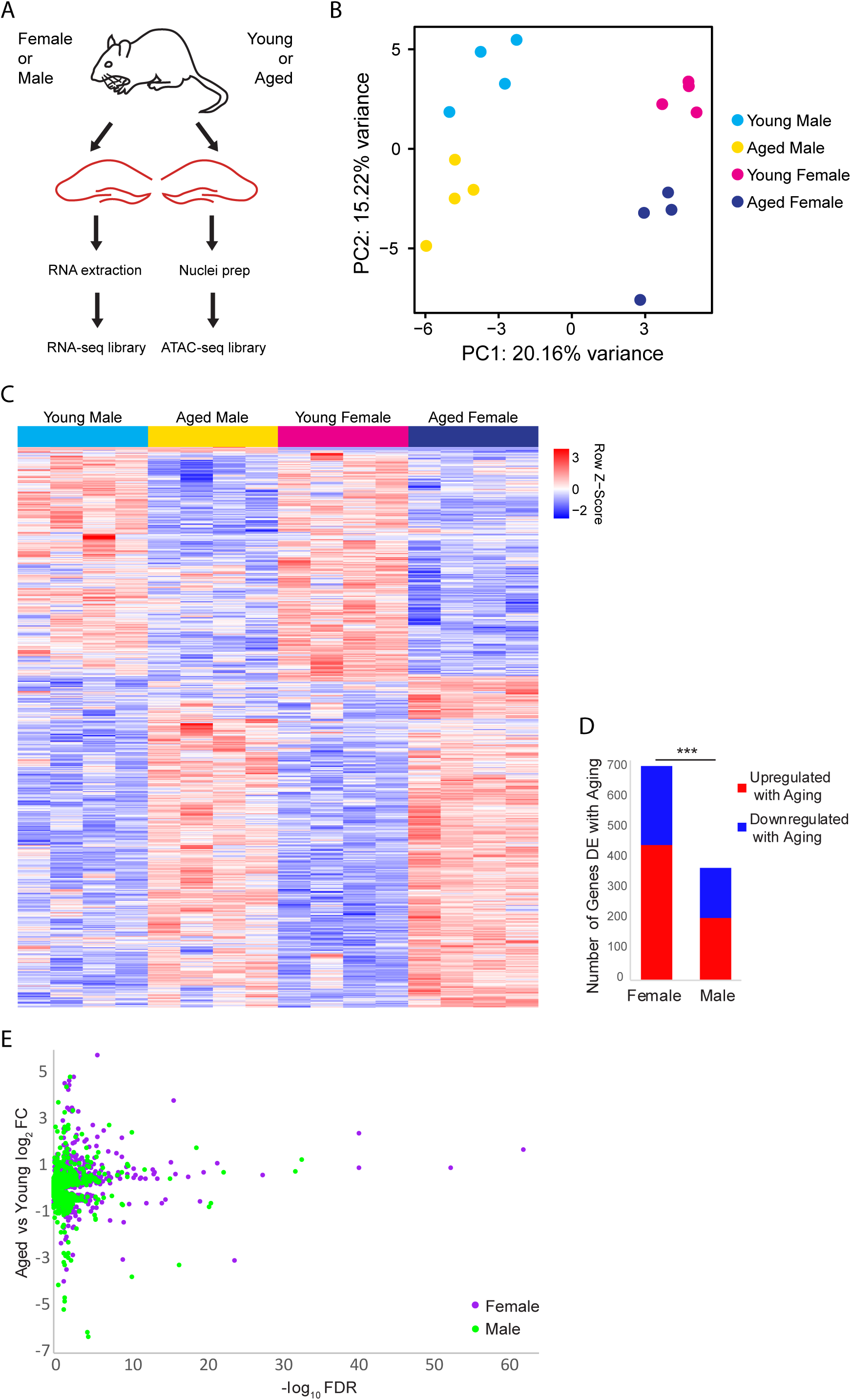
Gene expression changes with aging and sex-bias in female and male mouse hippocampus. (A) Illustration of the experimental design (B) Principle component analysis (PCA) of the RNA-seq gene expression data. N = four biological replicates for each group (young adult female, aged female, young adult male, aged male). (C) Expression heat map of 905 sex or aging related differentially expressed (DE) genes (FDR < 0.05). Each row represents a gene and each column is a biological replicate. Red row z-scores indicate high expression and blue indicates low expression (see Supp. Table 1 for all FPKM and log_2_FC values). (D) Number of genes that are DE between aged and young animals in female and male (FDR < 0.05), *** indicates p < 0.001 for chi-square test. (E) Volcano plot of aging-related DE genes in males (green) and females (purple).

**Table 1:**
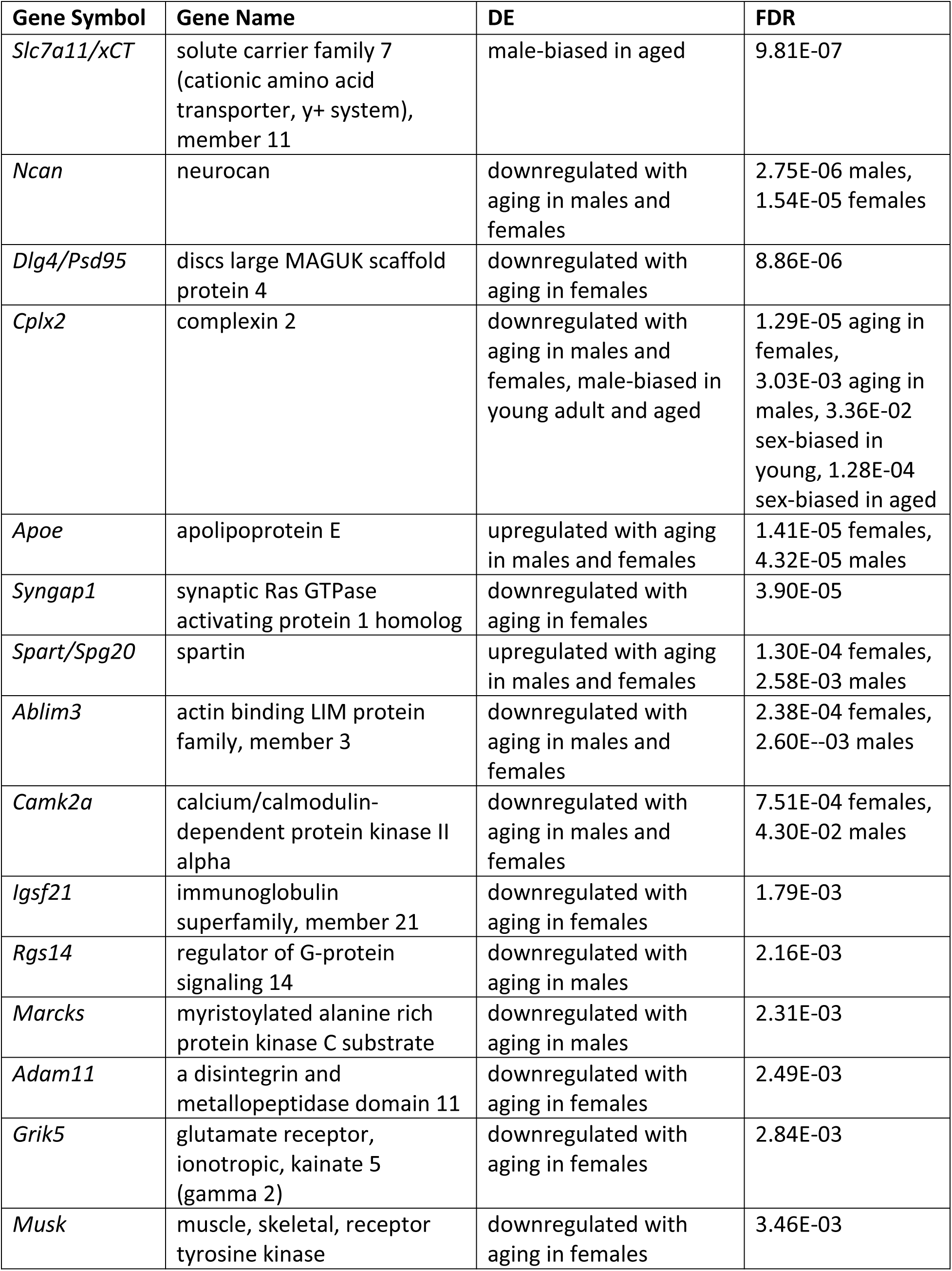
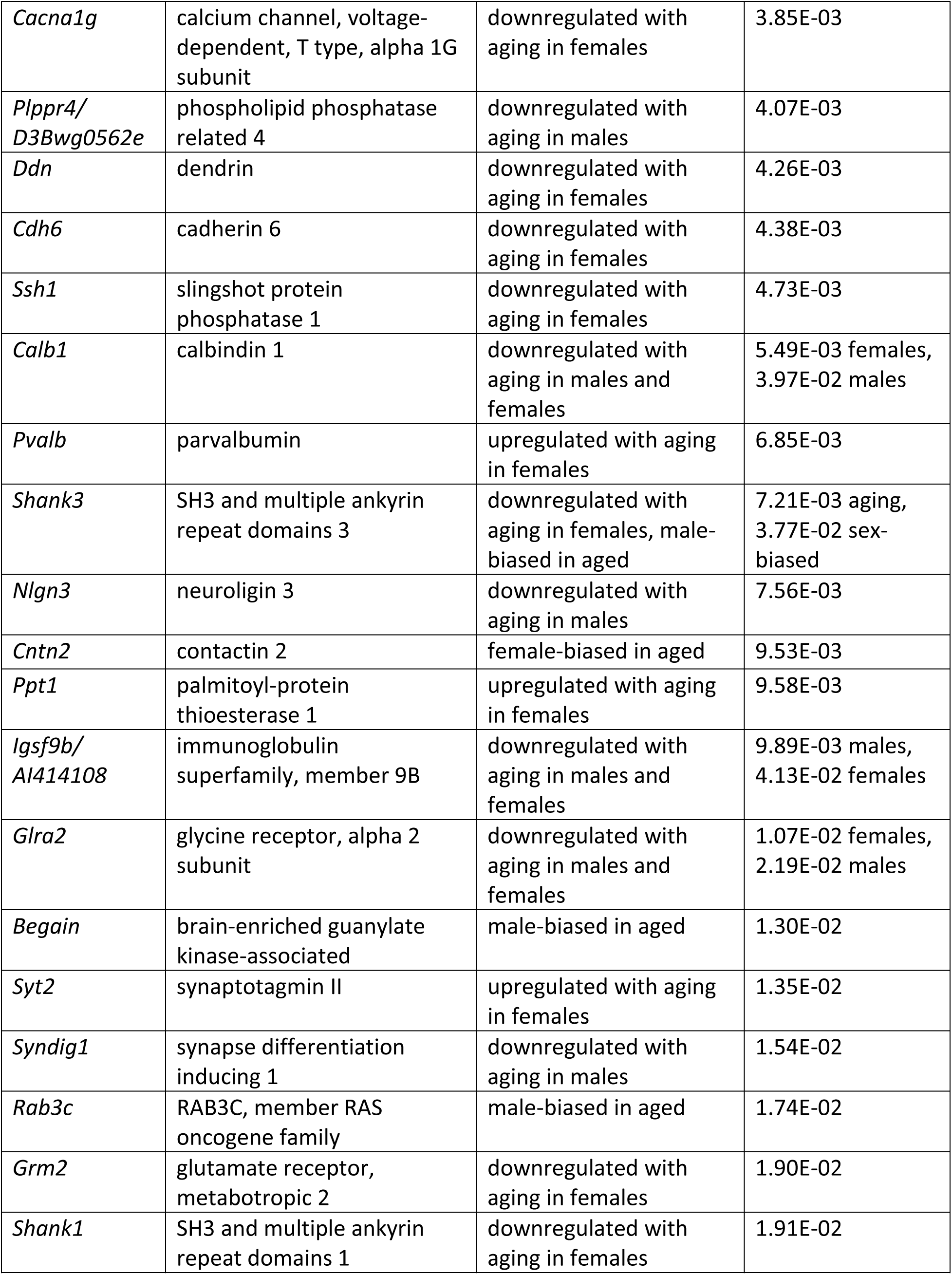
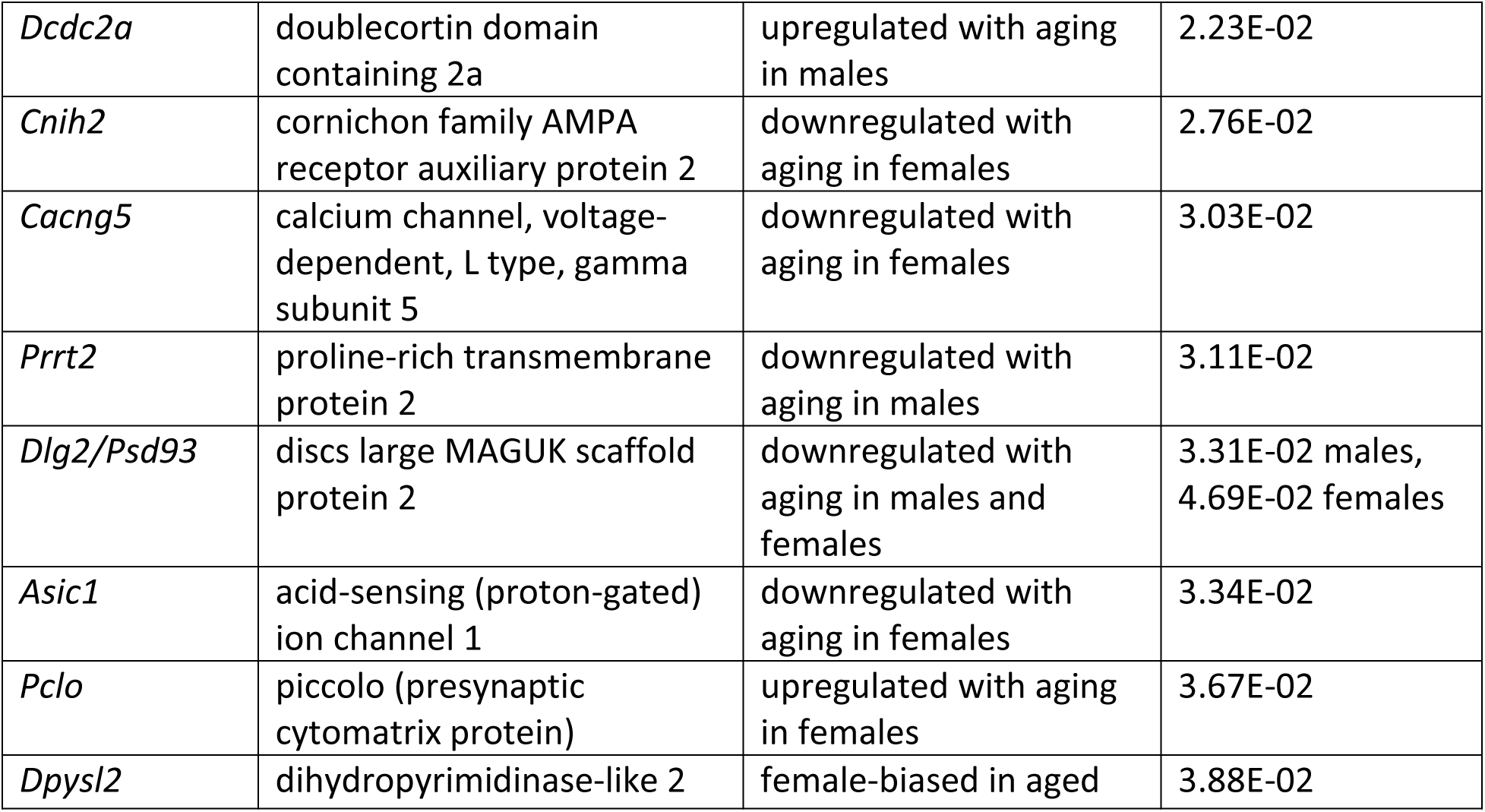
Synaptic-related DE genes. Hippocampal aging-related or sex-biased genes associated with synaptic function.

### More genes show sex-biased expression in the hippocampus with aging, including higher expression of myelin sheath-related genes in aged females compared to aged males

To determine whether and how sex impacts aging-dependent changes in gene expression, we compared gene expression profiles between males and females from young adult and aged hippocampus. In the hippocampus of young animals, we identified 17 sex-biased genes. Among these, neuropeptide Y (*Npy),* a neuropeptide whose expression is correlated with female-biased stress-related memory disorders^47^, showed a 1.3 fold higher expression in the hippocampus of young females compared to young males, but a female-specific downregulation with aging (young female vs male log_2_FC = 0.41, FDR < 0.001; average FPKM: young male = 64.34, aged male = 67.38, young female = 88.11, aged female = 71.34). In the hippocampus of aged animals, significantly more genes (42) were DE between sexes (chi-square test, p = 0.002, *X*^2^ = 9.8 Fig. 2A). Most of the sex-biased genes in the hippocampus of young animals also showed sex bias in the hippocampus of aged animals (Fig 2B). Of the genes with sex-biased expression in both young and aged, all were on sex chromosomes except for *Prl* (prolactin) and *Cplx2* (complexin 2). Complexin 2, a protein involved in synaptic vesicle fusion and whose dysregulated expression is associated with a number of cognitive disorders^48,49^, was more highly expressed in males compared to females in both age groups, and this gene was also significantly downregulated with aging in both sexes (female vs male: young log_2_FC = -0.16, aged log_2_FC = - 0.21; aged vs young: male log2FC = -0.16, female = -0.21). Of the sex-biased genes located on the X chromosome, all are known to escape X-inactivation in the mouse brain^50^. Interestingly, we also found that *Xist*, a female-specific noncoding RNA critical for X-inactivation^51,52^, was significantly upregulated in the female hippocampus with aging (log_2_FC = 0.21, FDR = 0.009; average FPKM: young male = 0.06, aged male = 0.04, young female = 27.75, aged female = 32.08). This finding is consistent with a recent report showing increased *Xist* expression with aging in mouse hypothalamus^10^. Altogether, these results suggest that aging results in more divergence in the hippocampal transcriptome between males and females.

**Figure 2:**
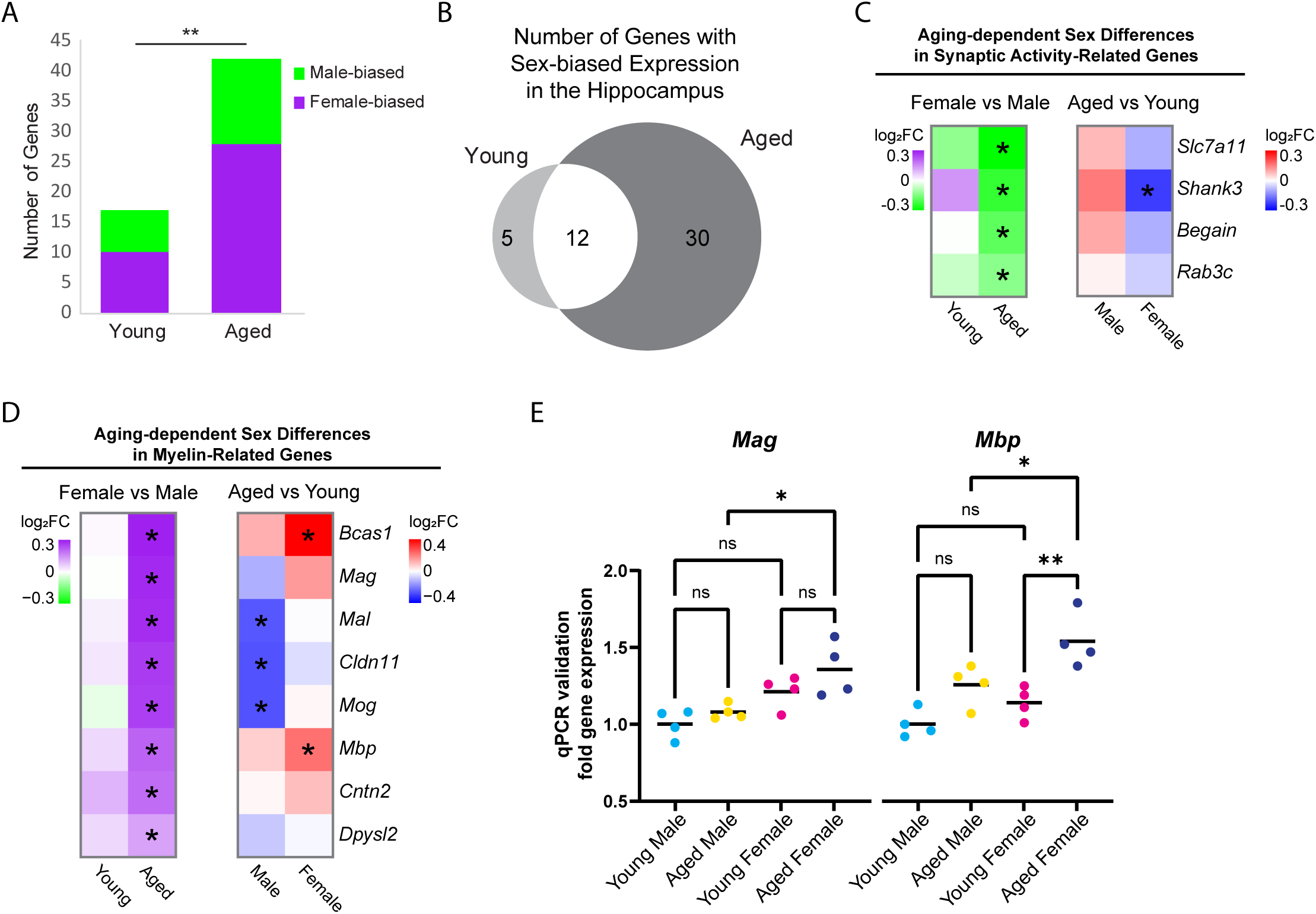
Sex differences in expression of hippocampal genes during aging. (A) Number of genes that were DE between female and male hippocampus (FDR < 0.05) in young adult and aged animals. Purple indicates higher expression in female vs male and green indicates higher expression in male vs female. ** indicates p < 0.01 by chi-square test. (B) Venn diagram of sex-related DE genes in the hippocampus. (C) Fold change heat maps of synaptic activity-related genes showing higher expression in aged male vs female hippocampus. Left heat map shows sex-related differential expression (purple indicates higher in female and green indicates higher in male). Right heat map shows aging-related differential expression.* indicates significant differential expression (FDR < 0.05). (D) Fold change heat maps (as in C) of myelin-related genes showing sex differences. (E) qPCR validation for myelin sheath genes *Mag* and *Mbp* in young and aged male and female hippocampus (n = four each condition). *Mag* aged male vs aged female ANOVA with Sidak’s multiple comparison test adjusted p = 0.022. *Mbp* aged male vs aged female ANOVA with Sidak’s multiple comparison test adjusted p = 0.039; young female vs aged female adjusted p = 0.004. * indicates p < 0.05, ** indicates p < 0.01, ns = not significant.

To further explore sex differences in gene expression that arise with aging, we focused on the 30 genes with sex-biased expression only in aged animals. Eight genes were male-biased (*Slc7a11/xCT, Begain*, *Rab3c*, *Shank3*, *Hist1h4d, D830046C22Rik*, *Ncl*, and *Scarf2*), the first four of which encode proteins involved in synaptic transmission (Fig. 2C)^53–56^. We found that 22 genes showed female-biased expression in aged hippocampus, including the myelination-related gene, *Bcas1,* which showed an aging-related increase in expression in female but not male hippocampus, resulting in significantly higher expression in aged females compared to aged males (log_2_FC = 0.36, FDR < 0.001; average FPKM: young male= 33.4; aged male = 37.3, young female = 33.8, aged female = 47.9).

Gene ontology analysis of female-biased genes in aged hippocampus revealed a significant enrichment of the term myelin sheath (GO:0043209; FDR < 0.001). We then assessed all genes associated with the GO terms “myelin sheath” and “myelination” (GO:0043209 and GO:0042552), and found that 8 genes (*Bcas1*, *Ma*g, *Mal*, *Mbp*, *Cldn11*, *Cntn2*, *Mog,* and *Dpysl2*) had sex-biased expression in the hippocampus of aged animals, all of which showed a female bias (Fig. 2D), whereas other oligodendrocyte-related genes such as *Olig1*, *Olig2*, *Opalin*, *Cnp*, *Plp1, Myrf, Gpr17* and *Mobp* were not DE between aged female and aged male hippocampus, suggesting that this was not due to a difference in the number of oligodendrocytes or oligodendrocyte precursor cells (OPCs) as has been previously reported^12^. Furthermore, we found no sex bias in the expression of myelination-related genes (GO:0043209 and GO:0042552) in young adult animals. The sex-bias in myelin genes in aged hippocampus resulted from aging-related changes in both males and females: aging resulted in a significant downregulation of *Mog*, *Cldn11* and *Mal* in the hippocampus of males but not females, whereas aging resulted in significant upregulation in *Bcas1* and *Mbp* in the hippocampus of females but not males (Fig. 2D). We also performed qPCR for two of the myelin sheath genes, *Mag* and *Mbp*, using samples from a separate cohort of animals and verified that the expression of both had female-biased expression in aged hippocampus (Fig. 2E). This sex-biased expression of myelin sheath genes suggests that the hippocampus of aged females may have less myelin degeneration or more remyelination than aged males, in agreement with previous studies reporting more remyelination and increased white matter volume in aged female versus aged male rats^57,58^.

### Aging results in changes in the expression of genes involved in cell adhesion, immune function and neuronal development in both sexes

Next, we focused on sex-independent gene expression changes in the hippocampus with aging. We found that 136 genes were upregulated and 64 genes were downregulated with aging in both sexes (Fig. 3A). We noted that although these genes were DE with aging in both sexes, for many genes the aging-related fold change in female hippocampus was larger than that in male hippocampus (Fig. 3B). We ranked DE genes by lowest FDR in either males or females and found that many of the aging-dependent genes (e.g. *C4b*, *Pcdhb9, Abca8a*, *Gfap*, *Igfbpl1*, *Zc3hav1*, *Ptpro*, *Gpr17* and *Il33*) have been previously reported to undergo similar age-related changes in the brain^8,11,12,59–61^(Fig. 3C). Gene ontology analysis revealed that genes upregulated with aging showed enrichment for cell adhesion (GO:0007155, FDR = 5.92E-08) and innate immune response (GO:0045087, FDR = 0.02), whereas genes downregulated with aging showed enrichment for nervous system development (GO:0007399, FDR = 5.38E-04) and neuron migration (GO:0001764, FDR = 0.002; Fig. 3D).

**Figure 3:**
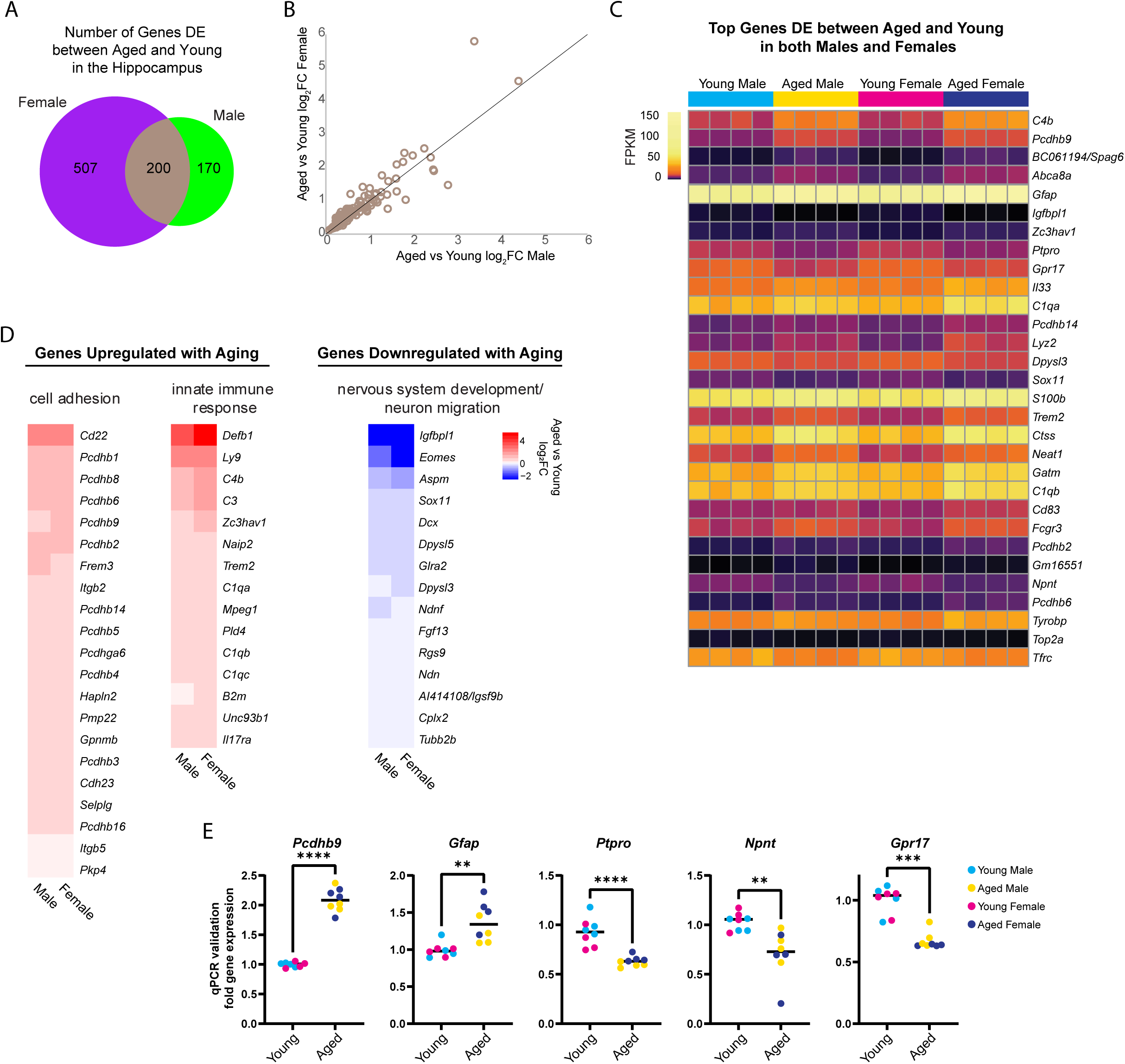
Aging is associated with changes in expression of cell adhesion, immune response and nervous system development genes. (A) Venn diagram of number of genes DE (FDR < 0.05) with aging in female and male mouse hippocampus. (B) Scatter plot of aged vs young adult log_2_FC in males and females for the 200 genes that showed aging-related differential expression in the hippocampus of both sexes. (C) Expression heat map of top 30 genes (sorted by lowest FDR) DE in both male and female hippocampus with aging. Each row represents a gene and each column is a biological replicate. FPKM for each gene/replicate is represented by color (see Supp. Table 1 for all FPKM and log_2_FC values). (D) Fold change heat maps of genes in biological process categories from gene ontology analysis that were upregulated (red) and downregulated (blue) in the hippocampus with aging. (E) qPCR validation for genes *Pcdhb9*, *Gfap*, *Ptpro*, *Npnt* and *Gpr17* in young and aged male and female hippocampus (n = four each sex/age). *Pcdhb9* unpaired t-test p < 0.0001; *Gfap* unpaired t-test p = 0.001; *Ptpro* unpaired t-test p < 0.0001; *Npnt* unpaired t-test p = 0.003; *Gpr17* Mann-Whitney test p < 0.001.). ** indicates p < 0.01, *** indicates p < 0.001, **** indicates p < 0.0001.

Increases in inflammatory and immune pathways are known to occur in the brain with aging, including increases in innate immune response genes in the hippocampus^11–13,62^. Consistent with these findings, we found many innate immune response genes that were upregulated with aging in both sexes (*B2m*, *C1qa*, *C1qb*, *C1qc*, *C3*, *C4b*, *Defb1*, *Ly9*, *Naip2*, *Trem2*, *Mpeg1, Pld4, Il17ra*, *Unc93b1*, *Zc3hav1*) (Fig. 3D), a group that includes three (*C4b*, *Zc3hav1, C1qa*) of the top 10 aging-upregulated genes.

Four of the most significantly aging-upregulated genes encoded protocadherins (*Pcdhb9*, *Pcdhb14*, *Pcdhb2*, *Pcdhga6*; Fig. 3C). Protocadherins are calcium-dependent cell adhesion molecules (CAMs) that are critical for dendrite morphology and synaptogenesis^63^. Within the class of cell adhesion genes that were upregulated with aging, half were protocadherin genes (*Pcdhb1*, *Pcdhb2*, *Pcdhb3*, *Pcdhb4*, *Pcdhb5*, *Pcdhb6*, *Pcdhb8*, *Pcdhb9*, *Pcdhb14*, *Pcdhb16*, *Pcdhga6*; Fig. 3D). To validate these findings, we performed qPCR for one of the protocadherin genes (*Pcdhb9*) using samples from a separate cohort of animals, which showed upregulation with aging (p < 0.0001; Fig. 3E).

Of the top most significant aging-downregulated genes, several are known to encode proteins that are located at or interact with the extracellular matrix (ECM; e.g. *Ptpro*, *Tfrc* and *Npnt*). We validated these findings by qPCR and found that *Ptpro* and *Npnt* decreased with aging in both males and females (*Ptpro* p < 0.0001, *Npnt* p < 0.01; Fig. 3E), whereas *Tfrc* decreased with aging in males but not females (data not shown). Given that CAMs and ECM proteins have been shown to be important for maintaining the integrity of synapses and synaptic plasticity^64,65^, these findings suggest that aging-related changes in the expression of both classes of genes in the hippocampus may affect synapse formation and signaling.

Aging resulted in the downregulation of genes in the hippocampus involved in nervous system development and neuron migration (*Igfbpl1, Eomes*, *Aspm, Dcx*, *Ndnf, Sox11, Dpysl3*, *Dpysl5*, *Fgf13*, *Glra2*, *Rgs9*, *Igsf9b, Ndn*, *Cplx2, Tubb2b;* Fig. 3D), including four of the most significantly downregulated genes (*Igfbpl1*, *Dpysl3*, *Sox11*, *Eomes*). We also found that *Gpr17,* a receptor critical for oligodendrocyte maturation^66,67^, was significantly downregulated with aging in both males and females (Fig. 3C). To validate these findings, we performed qPCR and detected a significant decrease in the expression of *Gpr17* in the hippocampus with aging (p< 0.001; Fig. 3E). These results are consistent with previous work showing a decrease in neurodevelopment-related genes and *Gpr17* in the brain with aging^68,69^.

Overall, we found many aging or sex-biased DE genes that are important for synaptic function (Table 1). Most of the DE synaptic-related genes showed an aging-related decrease in expression in either male or female hippocampus and encode proteins critical for the excitatory postsynaptic terminal such as DLG4/PSD95, DLG2/PSD93, SYNGAP1, SHANK1, SHANK3, CACNG5, CAMK2A and RGS14. A small number of synaptic-related genes showed increased expression with aging, including the apolipoprotein gene *Apoe* (in both sexes), the primary cilia gene *Dcdc2a* (in males) and the gene encoding the presynaptic protein SPART/SPG20 (in both sexes)^70–72^. Additionally, we found aging-related changes in expression of genes regulating inhibitory neuron synaptic functions, including decreased expression of the inhibitory synaptogenic gene *Igsf21* (in females), the inhibitory synaptic adhesion gene *Igsf9b* (in both sexes) and calcium-binding protein gene *Calb1* (in both sexes) and increased expression of the calcium-binding protein gene *Pvalb* (in females) and inhibitory synaptic vesicle gene *Syt2* (in females)^73,74^. Together, these findings support previous reports of changing excitatory and inhibitory synaptic dynamics in the hippocampus with aging^75,76^.

### Alternative splicing of myelin sheath-related genes changes with aging in both sexes

Alternative splicing patterns have been reported to be altered with aging in the mouse hippocampus and show sex-specific differences in the human brain^11,77,78^. To examine sex bias and aging-related changes in alternative splicing, we generated a poly-A selected RNA-library from the hippocampus of young adult and aged male and female mice and queried five types of alternative splicing (AS) events: skipped exon (SE), alternative 5’ splice site (A5SS), alternative 3’splice site (A3SS), mutually exclusive exons (MXE), and retained introns (RI). We identified 591 significant AS events in 452 genes that showed sex-bias or aging-related changes (FDR< 0.05, Supp. Table 2). Aging in male hippocampus resulted in 171 AS events in 154 genes, significantly more events than occurred with aging in female hippocampus (Fig. 4A; 125 AS events in 124 genes; chi-square test, *X^2^*= 7.2, p = 0.007).

**Figure 4:**
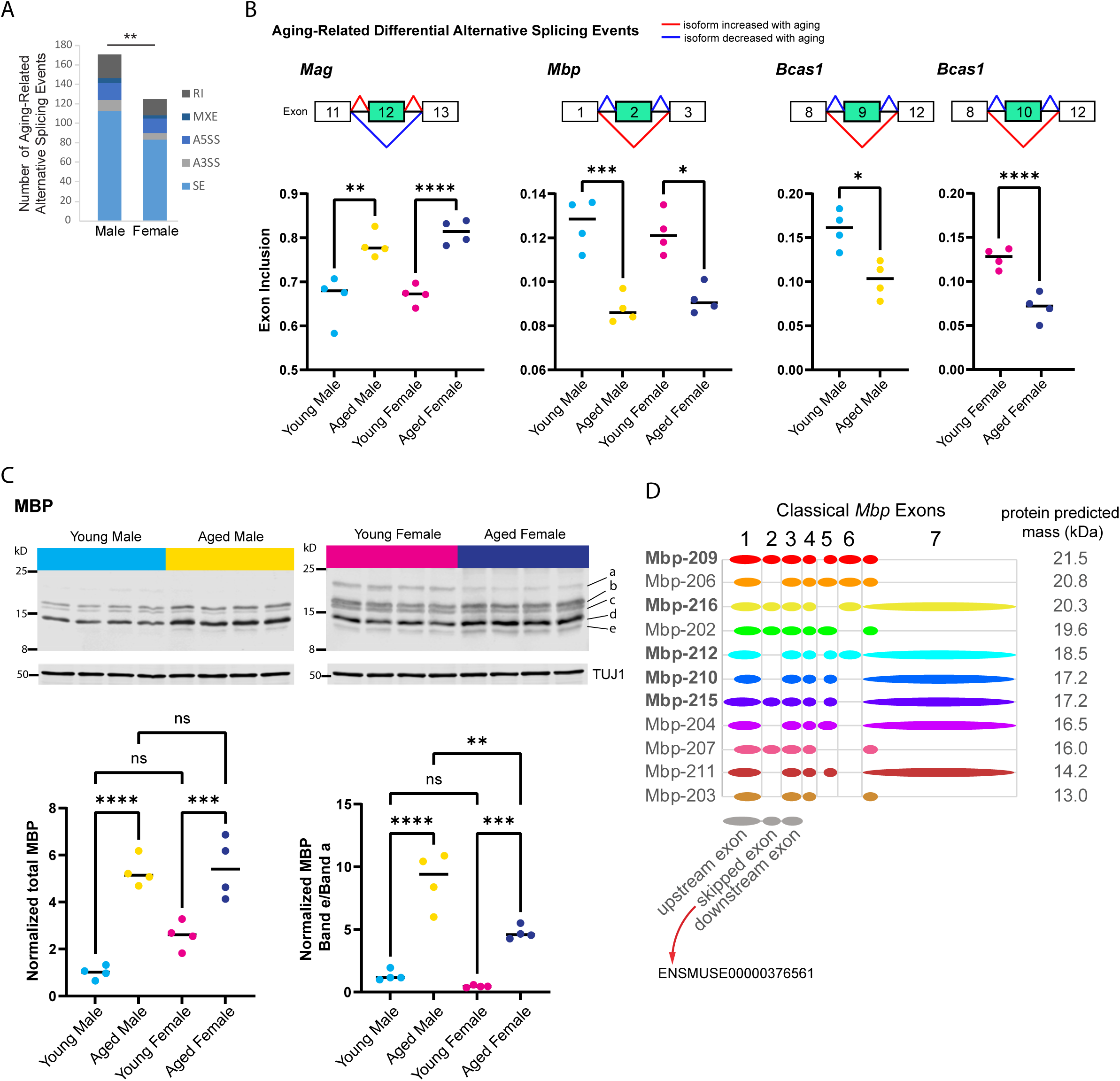
Aging is associated with alternative splicing of myelin sheath related genes. (A) Number of each type of aging-related AS event in male and female hippocampus. SE = skipped exon, A5SS = alternative 5’ splice site, A3SS = alternative 3’splice site, MXE = mutually exclusive exons, RI = retained intron. (B) Genes that exhibited aging-related alternative splicing events in both male and female hippocampus. Under each gene name is a diagram showing the skipped and flanking exons involved in each alternative splicing event. Below the diagram, plots show exon inclusion values and statistical significance calculated from rMATS^27^. *Mag* aging-related inclusion of exon 12 (ENSMUSE00000373997) in males rMATS FDR = 0.001 and females FDR < 0.0001. *Mbp* aging-related skipping of in exon 2 (ENSMUSE00000376561) in males FDR < 0.001 and females FDR = 0.016. *Bcas1* aging-related skipping of exons 9-11 (ENSMUSE00000170506, ENSMUSE00000170504, ENSMUSE00001234083) in males FDR = 0.029 and female FDR < 0.0001 (also see Supp. Fig1. A-C). (C) Western blot analysis for MBP protein isoforms in young adult and aged hippocampus. Top row shows western blot images for MBP protein and loading control TUJ1 (below) for young and aged male and female hippocampus (4 biological replicates each). Letters on right side indicate the bands that were quantified. Bottom panels show quantification of total MBP (left plot) and the ratio of the lowest-migrating MBP isoform (e) to the highest-migrating MBP isoform (a). See Supp. Fig. 1D-E for quantification of each band. Total MBP young male vs aged male ANOVA with Sidak’s multiple comparison test adjusted p < 0.0001; young female vs aged female adjusted p = 0.001. MBP band e/a young male vs aged male ANOVA with Sidak’s multiple comparison test adjusted p < 0.0001; young female vs aged female adjusted p = 0.001; aged male vs aged female adjusted p = 0.001.). (D) Classical (not including Golli) *Mbp* protein-coding splice variants from Ensembl^190^ release 102 with a diagram of included exons. Diagram of upstream, skipped and downstream exons are shown in gray below splice variant diagrams. Bolded transcript names indicate Ensembl’s designation as stable, reviewed, and high quality transcript annotations. The predicted protein mass from Ensembl for the product of each transcript is indicated to the right of the diagram. ** indicates p < 0.01, *** indicates p < 0.001, **** indicates p < 0.0001, ns = not significant.

**Table 2:**
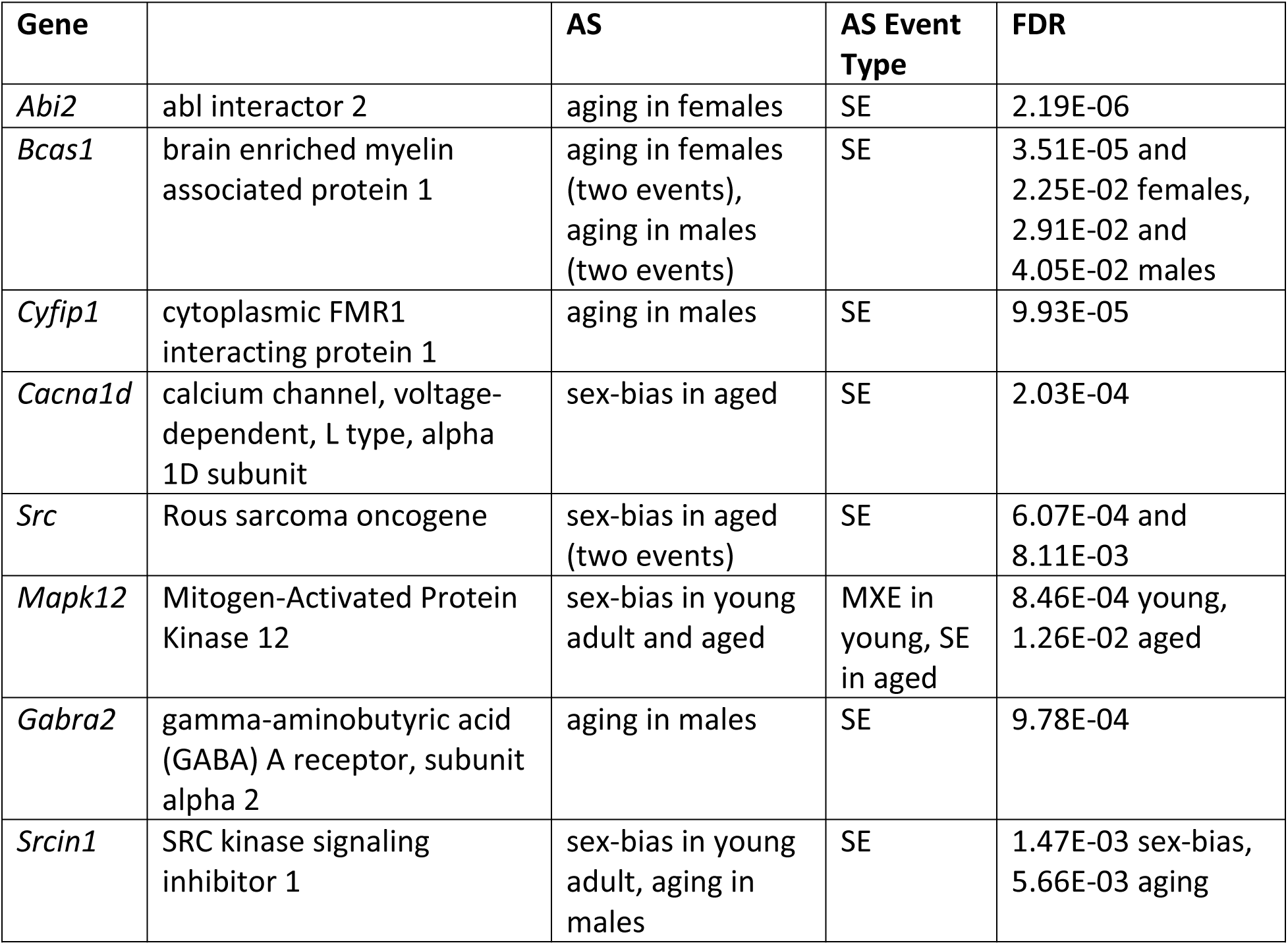
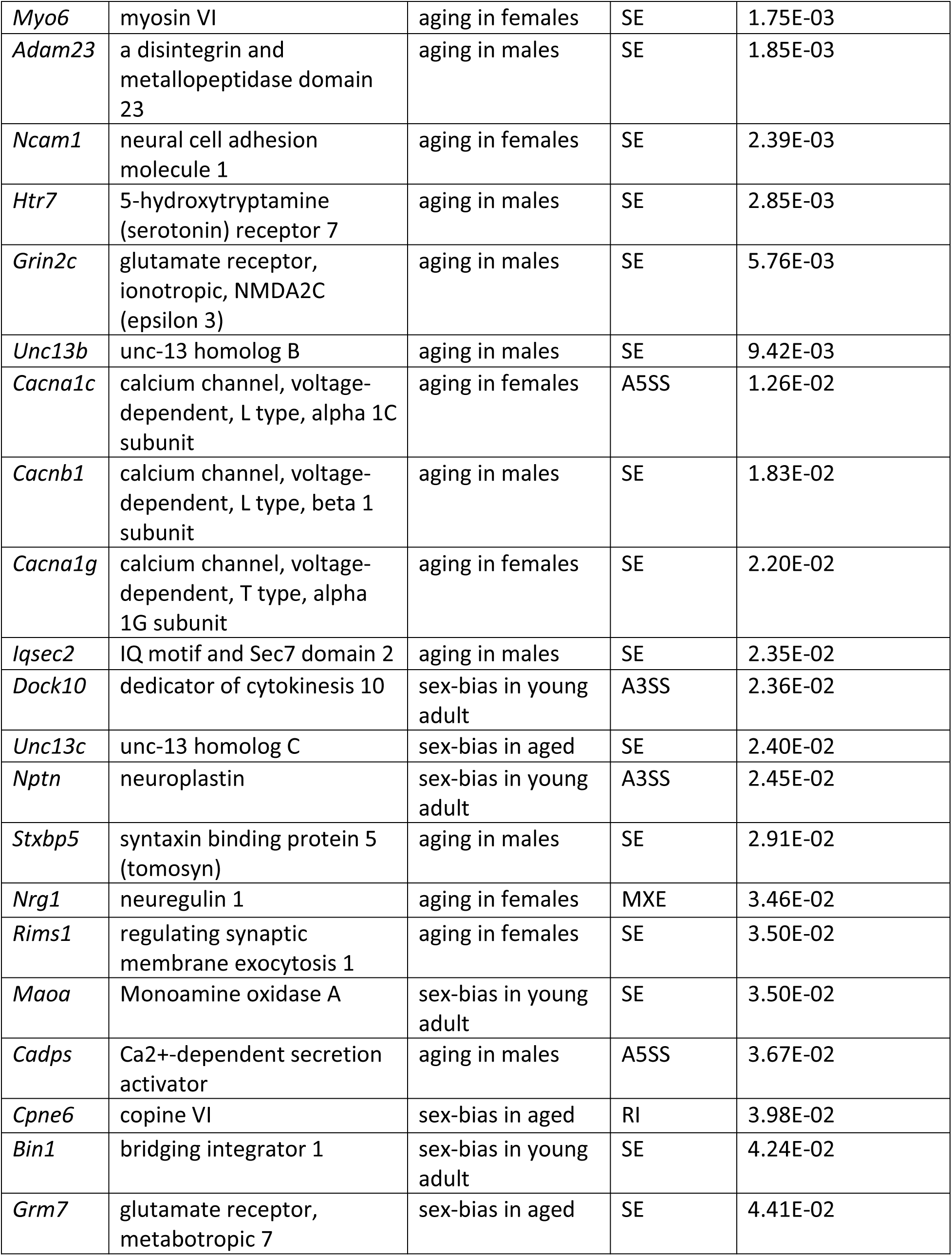
Synaptic-related AS genes. Alternative splicing events in the hippocampus in genes associated with synaptic function. SE = skipped exon, A5SS = alternative 5’ splice site, A3SS = alternative 3’splice site, MXE = mutually exclusive exons, RI = retained intron.

We focused on sex-independent AS in the hippocampus with aging, and identified seven genes (*Mag, Bcas1, Mbp, 4732471J01Rik, Huwe1, Ctxn3 and Mthfsl)* with skipped exon events that showed the same aging-related AS event in both males and females. The top three most significant aging-related AS events occurred in myelin-related genes: (1) inclusion of exon 12 (ENSMUSE00000373997) in *Mag*, (2) skipping of in exon 2 (ENSMUSE00000376561) in *Mbp*, and (3) skipping of exons 9-11 (ENSMUSE00000170506, ENSMUSE00000170504, ENSMUSE00001234083) in *Bcas1* (Fig. 4B, Supp. Fig 1A-C). As mentioned previously, the overall transcript abundance of myelin sheath genes *Mag*, *Mbp* and *Basc1* were female-biased in aged hippocampus (Fig. 2D), however the proportions of exon inclusion for these genes was similar between the sexes. The aging-related skipping of exon 2 in *Mbp* was also supported at the protein level: aged animals of both sexes exhibited an overall increase in MBP protein compared with young animals, but specifically showed a loss of a higher-migrating MBP species and a gain of a low-migrating species that correspond to *Mbp* splice variants with the inclusion or skipping of exon 2, respectively (Fig. 4C-D, Supp. Fig. 1D-E). These aging-related changes in *Mag* and *Mbp* exon skipping are similar to those that have been reported to occur throughout development and aging^79–83^. In addition to *Mag* and *Mbp*, *Bcas1* is also known to be involved in myelination^84,85^, therefore aging results in changes in the expression of different isoforms of myelin-related genes.

### Sex-biased alternative splicing in the hippocampus of young and aged hippocampus

We found 156 sex-biased AS events in 144 genes in young adult hippocampus and 139 sex-biased AS events in 129 genes in aged hippocampus (Fig. 5A), including 9 genes with sex-biased AS in both young and aged (*Ankrd13c, Ccdc62, Epn2, Lrrc14, Lrrcc1, Mapk12, Pofut2, Rtel1, Tle1*). We ranked the sex-biased AS events by lowest FDR-adjusted p-value in either young or aged hippocampus and found that within the 30 most significant autosomal AS events (Fig. 5B), many occurred in genes encoding nuclear proteins such as the polycomb repressive complex-2 protein EZH2, as well as other genes encoding DNA-binding proteins such as *Hmga1, Rorc, Ccdc62* and *Zmym5*. Furthermore, we detected an A5SS event that showed sex-biased AS in *Miat/Gomafu*, an lncRNA that regulates splicing, represses transcription through association with the polycomb repressive complex and is implicated in anxiety and schizophrenia^86,87^. One of top ten most significant AS events in young hippocampus was an A3SS splicing event in *Ift88*, a transport protein important for primary cilia maintenance in neurons^88^. Indeed, in young but not aged hippocampus, we found sex-biased AS in a number of genes associated with primary cilia including *Bbs4*, *Fuz*, *C2cd3*, *Rab34*, *Fam149b* and *Cdk10*.

**Figure 5:**
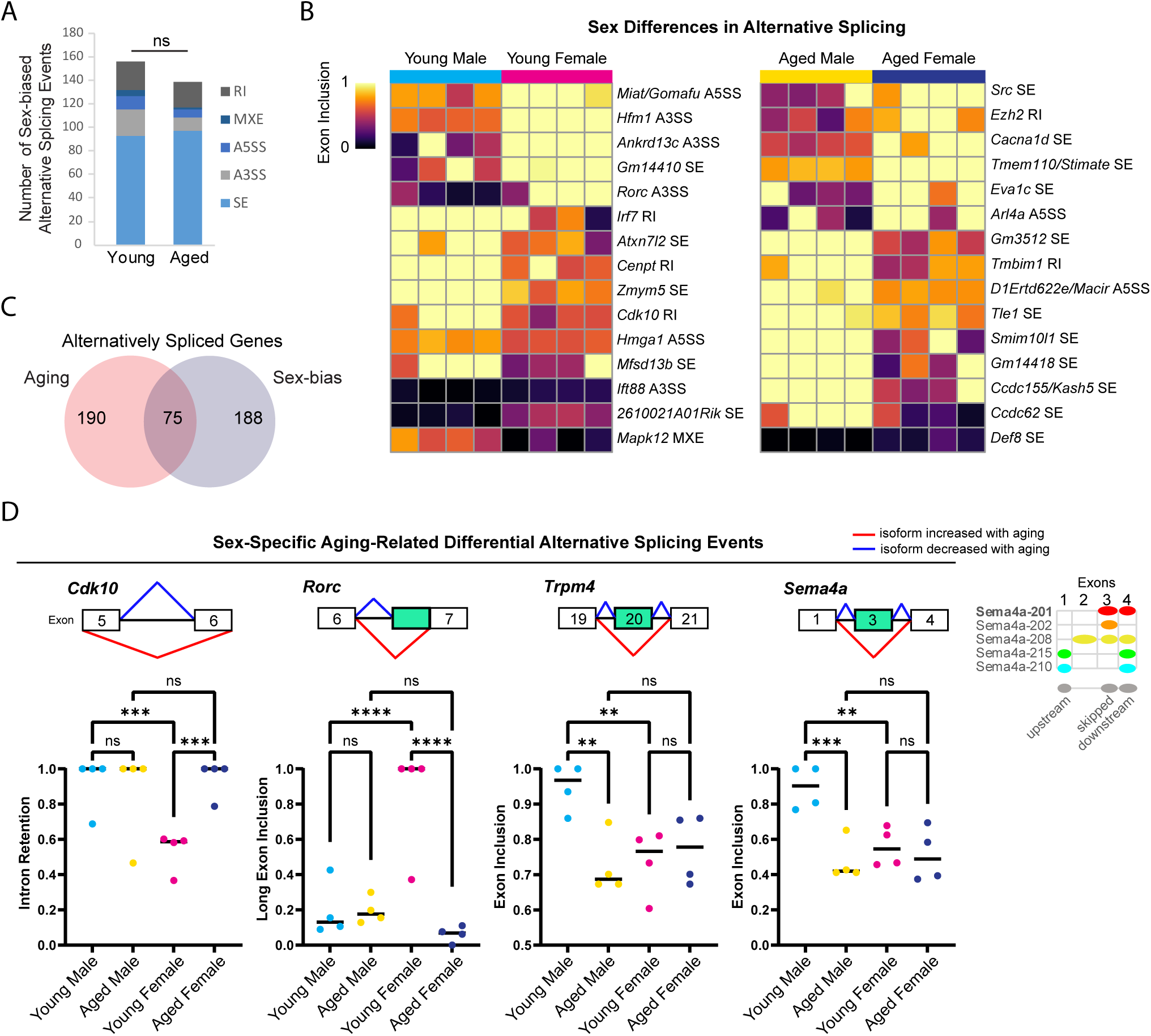
Sex differences in alternative splicing in the hippocampus. (A) Number of each type of sex-biased AS event in young adult and aged hippocampus. SE = skipped exon, A5SS = alternative 5’ splice site, A3SS = alternative 3’splice site, MXE = mutually exclusive exons, RI = retained intron. ns = not significant. (B) Exon inclusion heat map of top most significant sex-biased AS events in young or aged (by rMATS FDR). Each column represents a biological replicate, and each row represents an AS event, labeled on right. (C) Venn diagram showing the number of genes with aging-related or sex-biased AS events. (D) Example genes with sex-specific aging-related alternative splicing events in hippocampus. Under each gene name is a diagram showing the skipped and flanking exons involved in each alternative splicing event. Below the diagram, plots show exon inclusion values and statistical significance calculated from rMATS. *Cdk10* female-bias in exclusion of intron between exons 5 (ENSMUSE00000292260) and 6 (ENSMUSE00000292228) in young hippocampus (FDR < 0.001) and aging-related intron retention in females (FDR < 0.001). *Rorc* female-bias in long exon inclusion of exon 7 (ENSMUSE00000566508) in young hippocampus (FDR < 0.0001) and aging-related short exon usage in females (FDR < 0.0001). *Trpm4* male-bias in exon inclusion of exon 20 (ENSMUSE00000594904) in young hippocampus (FDR < 0.01) and aging-related exon skipping in males (FDR < 0.01). *Sema4a* male-bias in exon inclusion of exon 3 (ENSMUSE00000832345) in young hippocampus (FDR < 0.01) and aging-related exon skipping in males (FDR < 0.001). Diagram on right shows Ensembl protein-coding transcripts for *Sema4a* with the skipped exon corresponding to the second exon of Sema4a-208. ** indicates p < 0.01, *** indicates p < 0.001, **** indicates p < 0.0001, ns = not significant.

For genes on the X chromosome, we found sex-bias in 10 splicing events in nine genes. For example, in aged hippocampus, the X-chromosome gene *Huwe1*, encoding an E3 ubiquitin ligase linked to intellectual disability^89^, was associated with an SE event that showed a both a change with aging in females and a sex-bias in aged hippocampus. Overall, we found ∼30% of genes with sex-bias in AS, also showed a change in splicing with aging (Fig. 5C). Sex-specific aging-related alternative splicing events occurred in many of the genes mentioned above (e.g. *Ezh2*, *Rorc, Ift88*, *Fuz*, *Cdk10*) and included isoform ratios that were specific to either young male or young female hippocampus (Fig. 5D).

Overall, we found significant AS events in a number of genes important for synaptic function (Table 2). One of the most significant sex-biased AS events in aged hippocampus was female-biased inclusion of exon 8a *Cacna1d,* a gene important for synaptic plasticity that encodes the pore-forming subunit of an L-type voltage-gated Ca^2+^ channel. Mutations in *Cacna1d* exon 8a have been implicated in autism spectrum disorder^90–92^, and alternative splicing of exon 8 may contribute to channel voltage dependency differences^93^. We also found aging-related AS events in voltage-gated calcium channel genes *Cacna1c, Cacnb1* and *Cacna1g* that may impact calcium channel function^94–97^. We found that aging in male hippocampus resulted in changes in splicing of the NMDA receptor gene *Grin2c*, the GABA receptor *Gabra2* and the serotonin receptor gene *Htr7*. In addition, in male hippocampus we found an aging-related SE event in the postsynaptic gene *Cyfip1,* which is associated with autism spectrum disorder and schizophrenia^98^. We also detected sex-biased splicing (MXE in young, SE in aged) of *Mapk12*, a gene encoding the neuroprotective MAPK, p38γ, that is involved in regulating the post-synaptic density^99,100^.

### Aging increases chromatin accessibility in the hippocampus of males and females

Dynamic changes in chromatin accessibility are critical to the precise control of gene expression patterns have been shown to be altered with aging^16,19^. We used ATAC-seq to probe for chromatin accessibility changes during aging in the hippocampus of male and female mice. The hippocampus of young adult and aged males and females showed a similar number of ATAC peaks, with similar distribution of genome annotations: ∼40% intergenic, 35% introns, and 18% promoter-TSS (Fig. 6A, Supp. Fig. 2A). We analyzed a merged set of 92,233 replicated peaks (see Methods; Fig.6A, Supp. Fig. 2B) and, similar to the results from our RNA-seq analysis, principle component analysis of fragment counts from ATAC-seq regions showed that samples were separated by sex and age by the first and second principle component, respectively (Fig. 6B, Supp. Fig. 2C). Overall, 774 genomic regions were differentially accessible either with sex or aging (FDR < 0.05). There were 393 regions with sex-biased chromatin accessibility and 404 regions showing accessibility changes with aging (Fig. 6C), including 23 regions that showed changes with both aging and sex. More differentially accessible regions (DARs) exhibited aging-related changes in female compared to male hippocampus (366 versus 109 regions, chi-square test, *X^2^* = 139.4, p > 0.00001), and this result was consistent even when reduced numbers of female samples were included to match the number of male samples (Supp. Fig. 2D). Approximately 90% of aging-related DARs showed increased chromatin accessibility with aging, with intergenic regions making up the majority of DARs that either opened or closed with aging (Fig. 6D). For example, an intergenic region in chromosome 13 was ∼10 times more accessible in aged male and female hippocampus compared to young adult hippocampus (Fig. 6E left), whereas an intergenic region in chromosome 8 was ∼1.5 times less accessible in aged compared to young adult hippocampus (Fig. 6E right). On average, regions that opened with aging showed a 3.4 and 2.3-fold increase in accessibility with aging in males and females, respectively, and those regions that closed with aging showed a 1.8-fold aging-related decrease in accessibility in both males and females (Fig. 6F). Altogether, these results indicate that aging leads to increased chromatin accessibility in the hippocampus, especially in non-coding regions.

**Figure 6:**
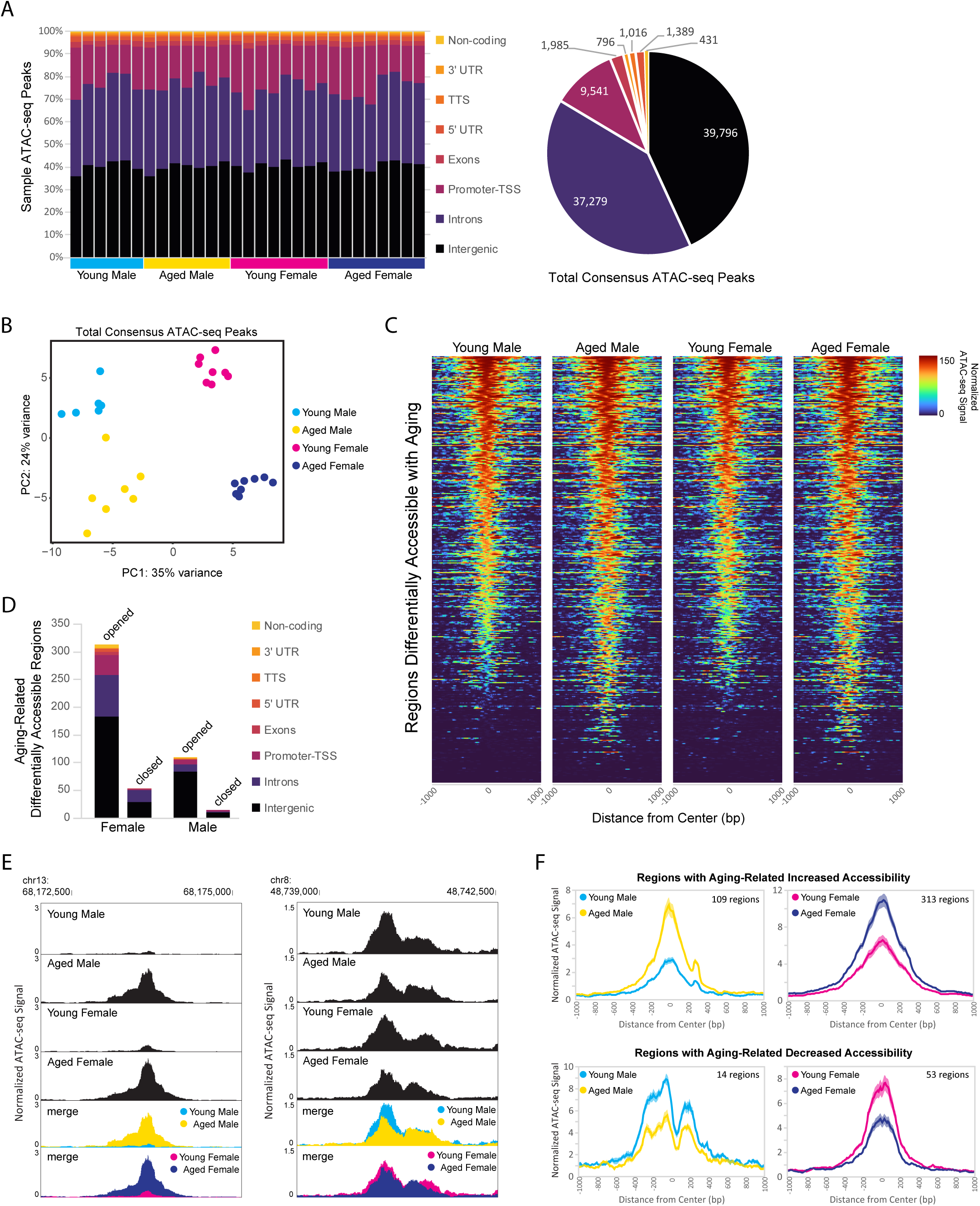
Aging is associated with higher chromatin accessibility, with most of the changes occurring in intergenic regions. (A) Genome annotation of individual sample ATAC-seq peaks, with each column representing a biological replicate (left) and genome annotation for total consensus ATAC-seq peaks (right). (B) Principle component analysis of ATAC-seq fragments in the total consensus peak set. (C) Heat maps of average normalized ATAC-seq signal per condition for aging-related differentially accessible regions (DARs). Normalized ATAC-seq signal/10 bp bin is shown for +/- 1000 bp around ATAC-seq region center, and regions are sorted by young male ATAC signal in the +/- 100 bp around center. (D) Genome annotation of regions that were differently accessible (FDR < 0.05) between young adult and aged hippocampus. (E) Browser track examples of regions showing aging-dependent differences in chromatin accessibility. Panel on left shows the average normalized ATAC-seq signal from young and aged male and female hippocampus in the intergenic-MuRRS-int|LTR|ERV1 region chr13: 68173077-68174082 that was significantly more accessible in aged compared to young males and females (male aged vs young log_2_FC = 3.85, FDR < 0.0001; aged vs young log_2_FC = 3.00, FDR < 0.0001). Panel on right shows the average normalized ATAC-seq signal in the intergenic region chr8: 48739982-48741266 that was significantly more accessible in young compared to aged males and females (male aged vs young log_2_FC = -0.63, FDR = 0.04; female aged vs young log_2_FC = -0.57, FDR =0.02). (F) ATAC-seq profiles for aging-related DARs in either males or females (as in E). Solid lines indicate the average of each condition’s normalized histogram (n = 6 young male, 7 aged male, 8 young female, 8 aged female) with shading indicating s.e.m.

Next, we focused on the sex-independent chromatin accessibility changes in the hippocampus with aging. We found that 85 regions showed differential ATAC-seq signal with aging in both sexes (Fig. 7A), and 99% of these regions exhibited aging-related increases in chromatin accessibility (84/85 regions opened, 1/85 closed). For the regions that gained accessibility in both sexes, there was a ∼3.8 fold increase in accessibility with aging (Fig. 7B), and motif enrichment analysis showed that these regions were enriched for HOXA and MEF2-like motifs (Supp. Fig. 3A). We ranked DARs by lowest FDR-adjusted p-value in either males or females and found that within the 30 most significant DARs, 13 were located in retrotransposable element-derived sequences, and all of these regions showed an opening in accessibility with aging (Fig. 7C). For example, an intergenic L1MA5A-type LINE-1 region in chromosome 14 showed a ∼4-fold increase in accessibility in the hippocampus of aged animals of both sexes compared to young animals (Fig. 7D). Twenty LINE-1 retrotransposon elements showed an aging-associated increase in accessibility in the hippocampus of both sexes (Supp. Fig. 3B). These regions encode truncated sequences incapable of retrotransposition; however, truncated LINE-1 sequences can be transcribed^101^, leading to DNA damage^102^ and immune activation^103,104^. To test for an aging-related increase in the transcription of LINE-1 elements in the hippocampus, we performed qPCR for the retrotransposon LINE-1 with three different primer sets. We found that with aging, there was an average increase of 32% in LINE-1 transcript abundance in the hippocampus (LINE-1 5’UTR p < 0.001, LINE-1 ORF1 p < 0.01, LINE-1 5’UTR:ORF1 p < 0.05; Fig. 7E), in agreement with the finding that LINE-ORF1 protein increased in the frontal cortex with aging^16^. Previous studies have found that LINE-1 retrotransposon activity is more common in neurons than other somatic cells and that it increases with aging^105–108^. We examined the chromatin accessibility of full-length intact LINE-1 genomic regions^41^ and found suppression of chromatin accessibility in these regions in both young adult and aged hippocampus (Supp. Fig. 3C), indicating that full-length LINE-1 regions as a whole remained repressed during aging. These results indicate that there is opening of chromatin during aging, especially in retrotransposon-derived sequences with an accompanying increase in LINE-1 transcript abundance, which may contribute to aging-related genome instability^107^.

**Figure 7:**
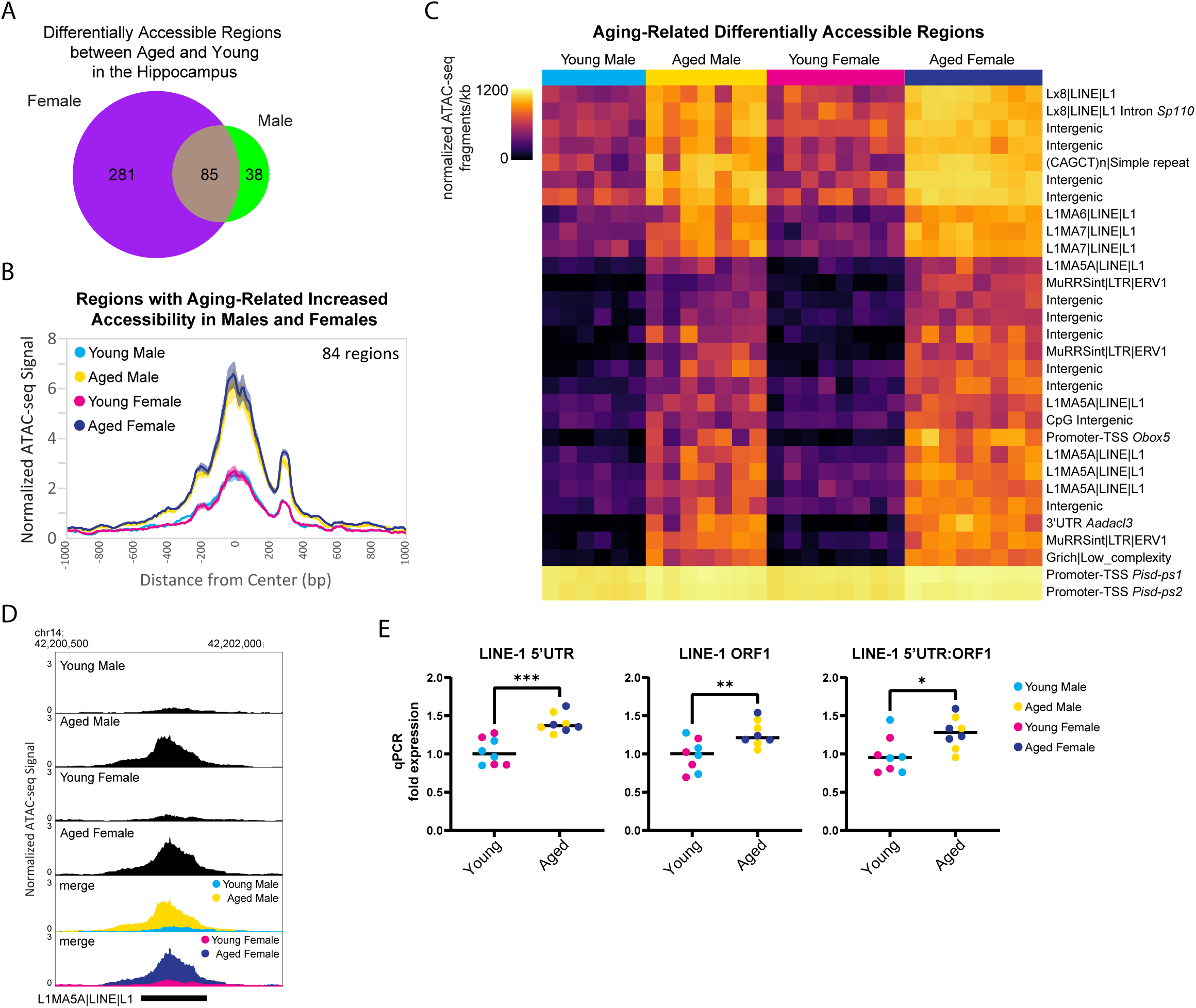
Aging is associated with higher accessibility in repetitive elements and higher expression of retrotransposon transcripts. (A) Venn diagram of number of DARs (FDR < 0.05) in young adult and aged hippocampus. (B) ATAC-seq profiles for regions that showed increased accessibility with aging in both males and females. Solid lines indicate the average of each condition’s normalized histogram (n = 6 young male, 7 aged male, 8 young female, 8 aged female) with shading indicating s.e.m. (C) Heat map of normalized total ATAC-seq signal per region for the top 30 aging-related DARs in both male and female hippocampus (top 30 by FDR, clustered by ATAC-seq signal). Each row represents a chromatin region with detailed annotation labeled on right, and each column is a biological replicate. Normalized ATAC-seq fragments/kb is represented by color (color scale is non-linear, see Supp. Table 3 for all normalized signal and log_2_FC values). (D) Browser track example of LINE-1 region showing aging-related increase in chromatin accessibility. Shown are the average normalized ATAC-seq signals from young and aged male and female hippocampus in the intergenic L1MA5A|LINE|L1 region chr14: 42200723-42201552 that was significantly more accessible with aging in males and females (aged vs young male log_2_FC = 2.22, FDR < 0.0001; aged vs young female log_2_FC = 2.05, FDR < 0.0001). (E) qPCR of LINE-1 transcript expression in young and aged hippocampus (n = four age/sex). LINE-1 5’UTR, LINE-1 ORF1 and LINE-1 5’UTR:ORF1 are three different primers for LINE-1 annealing to sequences within the 5’UTR, within ORF1, and spanning the 5’UTR and ORF1, respectively. Unpaired t-tests: LINE-1 5’UTR p < 0.001, LINE-1 ORF1 p = 0.009, LINE-1 5’UTR:ORF1 p = 0.02.).* indicates p < 0.05, ** indicates p < 0.01, *** indicates p < 0.001.

### Sex differences in chromatin accessibility include male-bias in promoter accessibility

To assess sex-dependent differences in chromatin accessibility in the hippocampus, we analyzed the DARs between males and females from young adult and aged mice. We identified 29 autosomal regions in young animals and 281 autosomal regions in aged animals that showed sex-biased differential accessibility, with aging associated with more sex differences in autosome chromatin accessibility (chi-square test, young vs aged *X^2^* = 205.2, p > 0.00001; Fig. 8A). Sex bias in chromatin accessibility resulted in ∼1.7 fold change in peak ATAC-seq signal in sex-biased DARs (Fig. 8B), relatively smaller than those seen with aging-related differential accessibility (Fig. 6F). When we ranked sex-biased regions by lowest FDR in either young or aged animals, we found that the majority of the most significant male-biased regions were annotated to promoters, whereas the most significant female-biased regions were intergenic (Fig. 8C). Similarly, when we analyzed the genomic ontology for all sex-biased DARs, we found that most regions that were more open in females versus males were annotated to intergenic regions (Fig. 8D). In contrast, most male-biased accessible regions were promoter-TSS regions (8/15 in young; 80/157 in aged). For example, an intergenic region (1.5 Mb from nearest gene, *Ncam2*) was more accessible in aged females compared to aged males (Fig. 8E left), and the TSS-promoter region of *Stub1* was more accessible in aged males compared to aged females (Fig. 8E right). Motif analysis revealed that promoters with male-biased accessibility were enriched for CpG island-associated motifs including CGCG/GFX and GC-box/SP/KLF motifs (Supp. Fig. 4A). Non-promoter regions with male-biased accessibility were enriched in CTCF, HOXA, ZNF382 and MEF2 motifs whereas non-promoter regions with female-biased accessibility were enriched for CTCF, E-box/NEUROD1, AP-1 and TLX motifs (Supp. Fig. 4B). Most mammalian promoter sequences are GC-rich and contain CpG islands^109^, and indeed we found the CpG content of accessible promoter regions in the hippocampus was ∼7% whereas that of accessible non-promoter regions was ∼2% (Supp. Fig. 5A,B). However, we found that in young and aged hippocampus, the CpG content in male-biased non-promoter regions was higher than average (∼4%; Kruskal-Wallis tests with Dunn’s correction, p < 0.05; Supp. Fig. 5B). To test if the sex differences in accessibility for promoter and CpG-rich regions could be due to GC bias, we used smooth-quantile with GC-content normalization^42,43^, which can correct for GC-content effects in ATAC-seq datasets. With this normalization technique, we found similar male-bias in accessibility at promoters (QSmooth-GC; Supp. Fig. 5C) and CpG-rich regions (Supp. Fig. 5A,B; p < 0.0001); therefore, although we cannot rule out GC-bias effects, these sex differences in accessibility appear to be biological in nature. Importantly, male-biased promoter accessibility did not appear to result in increased transcription at those genes, as none of the autosomal genes with male-biased accessibility at promoters showed sex-biased expression. Therefore, male and female hippocampus may compensate for sex differences in promoter accessibility through different mechanisms to regulate expression.

**Figure 8:**
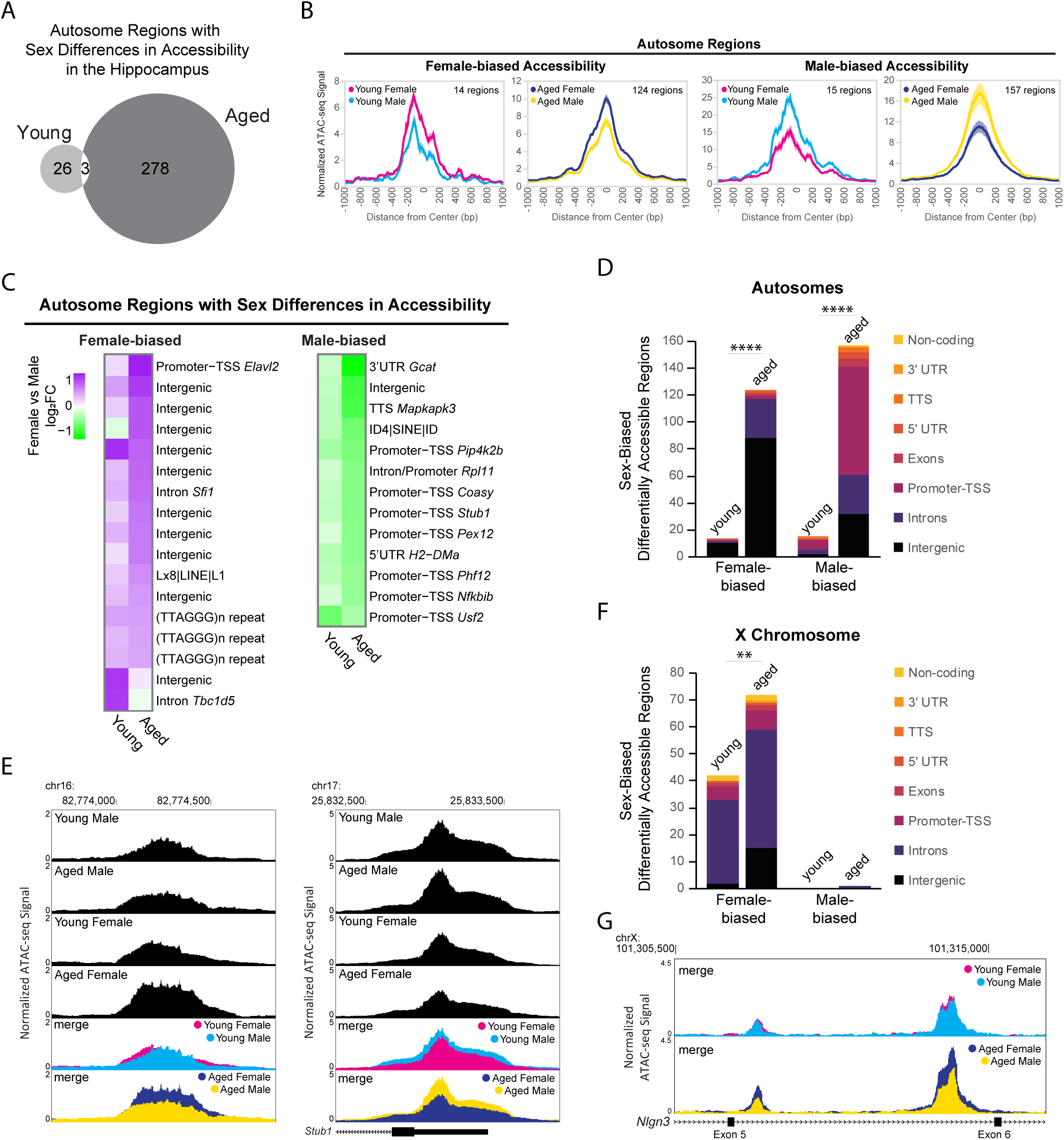
Sex differences in chromatin accessibility. (A) Venn diagram of number of autosome sex-biased DARs in young adult and aged hippocampus (FDR < 0.05). (B) ATAC-seq profiles for sex-related DARs on autosomes. Solid lines indicate the average of each condition’s normalized histogram (n = 6 young adult male, 7 aged male, 8 young adult female, 8 aged female) with shading indicating s.e.m. (C) Fold change heat map of sex-related differential accessibility for the top 30 autosome regions with differential accessibility between male and female hippocampus (top 30 by FDR, clustered by ATAC-seq signal; purple indicates higher accessibility in females and green indicates higher accessibility in males). Each row represents a chromatin region with detailed annotation labeled on right. (D) Genome annotation of autosome regions that were differently accessible between female and male hippocampus. Chi-square test, young vs aged *X^2^* = 87.7 more open female, *X^2^* = 117.3 more open male, p > 0.00001 (indicated with ****). (E) Browser track examples of regions showing sex differences in chromatin accessibility. Panel on left shows the average normalized ATAC-seq signal from young and aged male and female hippocampus in the intergenic region chr16: 82773954-82774686 (1.5 Mb from nearest gene, *Ncam2*) that was significantly more accessible in aged females than males (FDR = 0.003, female vs male log_2_FC = 0.79; one of two regions annotated to *Ncam2* showing female-bias in accessibility). Panel on right shows the average normalized ATAC-seq signal in the promoter-TSS region chr17:25832563-25833564 of *Stub1* that was significantly more accessible in aged males than females (FDR = 0.002, female vs male log_2_FC = -0.61). F) Genome annotation of chromosome X regions that were differently accessible between female and male hippocampus. Chi-square test, young vs aged *X^2^* = 8.1 more open female, p > 0.01 (indicated with **). (G) Browser track example of X chromosome region showing sex differences in chromatin accessibility. The average normalized ATAC-seq signal is shown from young and aged male and female hippocampus in the intronic region chrX: 101308500-101316000 between exons 5 (ENSMUSE00001248461) and 6 (ENSMUSE00001264564) of *Nlgn3*. Two regions were significantly more accessible in aged females than males, with the second ATAC-seq peak also showing female-specific opening with aging (region 101309331-101309581 aged female versus male log_2_FC = 0.88, FDR = 0.001; region 101313685-101314599 aged female vs male log_2_FC = 0.85, FDR =1.19E-11; female aged vs young log_2_FC = 0.70, FDR =1.11E-7).

We evaluated sex differences in chromatin accessibility of the X chromosome in young adult and aged hippocampus. Of the 2,595 ATAC-seq regions on the X chromosome, 74 regions (3%) showed sex differences in accessibility, with all but one region showing female-biased accessibility. We found 41 DARs on the X chromosome that showed a sex-bias in both young and aged animals, all of which showed female-biased accessibility. Most of these regions were located within *Dxz4* microsatellite and *Firre* locus important for X-chromosome 3D structure (29 female-biased regions in introns of *4933407K13Rik*, *Firre* and *Gm35612/CrossFirre*). Many other female-biased regions were located in the gene body or promoter of female-biased genes (*Kdm6a, Ddx3x, Eif2s3x, Kdm5c, Xist*) and/or genes located in the X inactivation center (*Xist*, *Ftx, Mir421, Jpx, Gm9159*)^110^. The number of regions with female-biased accessibility increased with aging (Fig. 8F; chi-square test, young vs aged *X^2^* = 8.1, p > 0.01), including aging-dependent female bias in accessibility in regions annotated to female-biased genes (*Kdm6a*, *Eif2s3x*, *5530601H04Rik*) as well as in the gene bodies of the synaptic regulators *Nlgn3* and *Srpx2*. For example, two regions in an intron of *Nlgn3*, a gene that encodes the synaptic protein neuroligin-3^111^ and whose expression was downregulated with aging in males, showed no sex bias in chromatin accessibility in young animals. However, a female-specific opening with aging resulted in female-biased accessibility in aged hippocampus (Fig. 8G). In young hippocampus, all X chromosome regions showed a female-bias in accessibility. Similarly, in aged hippocampus, all X chromosome regions were more accessible in females compared to males, except for one region that showed a male-bias in accessibility. This region spans the promoter, first exon and intron of *Gm35612/CrossFirre,* a lncRNA that is repressed by expression of *Firre,* which is a lncRNA involved in maintenance of X chromosome inactivation^112^. Altogether, these findings suggest that CpG-rich regions, including promoters, are more accessible in young and aged male compared to female hippocampus, and that there is more divergence in chromatin accessibility in the hippocampus between males and females with aging.

### Aging and sex-biased differential expression is correlated with chromatin accessibility in associated regions and TSSs

Transcription is a dynamic process involving the controlled coordination of regulatory elements such as promoters and enhancers, and DNA accessibility can indicate active regulatory elements bound by or permissive to chromatin binding factors^113,114^. Therefore, we wanted to assess the relationship between aging- or sex-related changes in chromatin accessibility and gene expression. For example, many of the genes in the *Pcdhb* gene cluster were upregulated with aging, and a corresponding increase in chromatin accessibility can be observed in the ATAC-seq signal in this region (Fig. 9A). Similarly, *Lin28b*, a gene encoding a RNA-binding protein that regulates miRNA maturation was upregulated with aging in the hippocampus of males and females, and a region upstream of *Lin28b* showed an aging-related increase in accessibility in the hippocampus of both sexes (RNA FDR < 0.001 and ATAC peak on far right, upstream/alternative promoter of *Lin28b,* FDR < 0.01 for both sexes; Fig. 9B).

**Figure 9:**
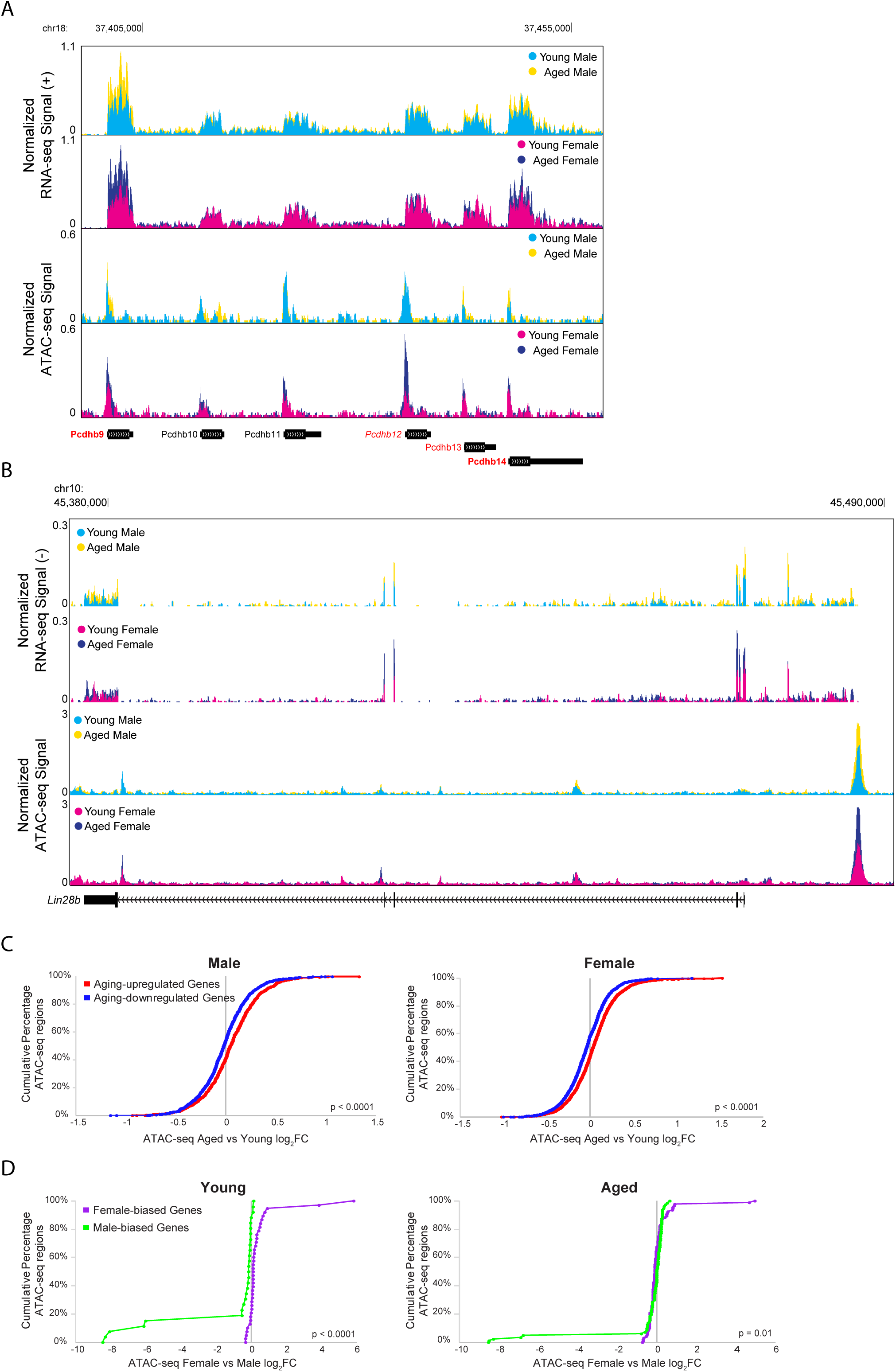
Aging and sex-related differential expression is correlated with ATAC-seq differential accessibility. (A) Browser track example showing RNA-seq and ATAC-seq signal for a portion of the *Pcdhb* gene cluster (region chr18: 37,397,808-37,458,667), containing four genes that were significantly upregulated with aging. The top two tracks show the normalized +-strand RNA-seq signal for male (first row) and female (second row) hippocampus. The bottom two tracks show the normalized ATAC-seq signal for male (third row) and female (fourth row) hippocampus. Genes that were upregulated with aging in both sexes are indicated with bold red (*Pcdhb9*: male aged vs young adult log_2_FC = 0.82, FDR-adjusted p = 7.21E-33, female aged vs young adult log_2_FC = 0.96, FDR-adjusted p = 1.68E-53; *Pcdhb14*: male aged vs young adult log_2_FC =0.46, FDR-adjusted p = 3.64E-6, female aged vs young adult log_2_FC = 0.68, FDR-adjusted p = 6.29E-17). One gene was upregulated with aging in female hippocampus only, indicated with italic red (*Pcdhb12*: male aged vs young log_2_FC = 0.23, FDR-adjusted p = 0.24, female aged vs young log_2_FC = 0.30, FDR-adjusted p = 0.02). One gene was upregulated with aging in male hippocampus only, indicated with regular font red (*Pcdhb13*: male aged vs young log_2_FC = 0.43, FDR-adjusted p = 0.006, female aged vs young log_2_FC = 0.28, FDR-adjusted p = 0.07). Genes that were not DE with aging are indicated with black font. (B) Browser track example as in A, showing RNA-seq and ATAC-seq signal for the *Lin28b* locus (chr10: 45,375,000-45,491,000). RNA signal suggests a long *Lin28b* isoform including additional exons present in the ENSMUST00000214555.1/Lin28b-202 transcript from an alternative promoter located at ATAC-seq peak. RNA upregulated with aging female: log_2_FC = 0.86, p =4.9E-7; male: log_2_FC 0.82, p = 8.3E-5; ATAC opened with aging female: log_2_FC = 0.60, p = 2.2E-3; male: log_2_FC = 0.75, p = 2.2E-4. (C) Cumulative distributions of aging-related chromatin accessibility fold changes for male (left panel) and female (right panel) hippocampus for ATAC-seq regions annotated to DE genes that were either upregulated (red) or downregulated (blue) with aging (male: Kolmogorov-Smirnov D = 0.14, p < 0.0001; female: Kolmogorov-Smirnov D = 0.15, p < 0.0001). (D) Cumulative distributions of sex-related chromatin accessibility fold changes for young (left panel) and aged (right panel) hippocampus for ATAC-seq regions annotated to DE genes that were either higher expressed in female (purple) or higher expressed in male (green) hippocampus (young: Kolmogorov-Smirnov D = 0.68, p < 0.0001; aged: Kolmogorov-Smirnov D = 0.24, p = 0.01).

To compare RNA-seq and ATAC-seq results across all DE genes, we first assessed the relationship between aging- or sex-bias fold changes in expression with the fold changes in chromatin accessibility for ATAC-seq regions closest to the TSS of those genes. We found that aging-related changes in gene expression were significantly correlated with aging-related changes in chromatin accessibility in male and female hippocampus (male: of 370 DE genes, 316 genes were associated with 2,044 ATAC-seq regions, Spearman r_s_ = 0.12, p < 0.0001; female: of 707 DE genes, 596 were associated with 3,600 regions, Spearman r_s_ = 0.15, p < 0.0001), and a stronger correlation was observed when including only ATAC-seq regions that were within 10 kb of the TSS (male Spearman r_s_ = 0.26, p < 0.0001, female Spearman r_s_ = 0.27, p < 0.0001). Similarly, ATAC-seq regions closest to the TSS of genes that were upregulated with aging in either male or female hippocampus showed more aging-related opening compared to ATAC-seq regions closest to the TSS of genes that were downregulated with aging (Fig. 9C; male: Kolmogorov-Smirnov D = 0.14, p < 0.0001; female: Kolmogorov-Smirnov D = 0.15, p < 0.0001).

We also found a correlation between sex bias in expression and chromatin accessibility in young hippocampus (young: of 17 DE genes, 16 were associated with 66 ATAC-seq regions, Spearman r_s_ = 0.74, p < 0.0001; aged: of 42 genes, 34 were associated with 178 ATAC-seq regions, Spearman r_s_ = 0.00, p = 0.96), and we found significant correlations in both young and aged hippocampus for regions within 10 kb of the TSS (young Spearman r_s_ = 0.88, p < 0.0001, aged Spearman r_s_ = 0.42, p < 0.01). Furthermore, ATAC-seq regions closest to the TSS of female versus male-biased genes showed higher chromatin accessibility in female versus male hippocampus in both young and aged animals (Fig. 9D; young: Kolmogorov-Smirnov D = 0.68, p < 0.0001; aged: Kolmogorov-Smirnov D = 0.24, p = 0.01).

To evaluate the relationship of aging and sex-bias DE with chromatin accessibility independent of consensus peaks, we analyzed the ATAC-seq signal surrounding DE genes’ TSSs. We found that chromatin accessibility increased in the region surrounding the TSS of genes that were upregulated with aging in female hippocampus (Supp. Fig. 6A; female: Kolmogorov-Smirnov p = 0.03, male: p = 0.11). Furthermore, chromatin accessibility was higher in female hippocampus in regions surrounding the TSS of female-biased genes, and chromatin accessibility was higher in males in regions surrounding the TSS of male-biased genes (Supp. Fig. 6B; female-biased expression, young: Kolmogorov-Smirnov p = 0.003; aged: p < 0.0001; male-biased, young: p < 0.0001; aged: p = 0.0001). In summary, we found that changes in expression tended to correlate with changes in chromatin accessibility, although we note that these correlations may be underestimated, as enhancers often do not regulate the gene with the nearest TSS^115,116^.

## Discussion

Memory impairments are a hallmark of aging. In both humans and rodents, the hippocampus is important for long-term memory formation^7,15,117^, and like humans, mice show memory impairments and changes in hippocampal function with aging^11,118–121^. While cell type abundance in the brain, including the hippocampus, does not appear to change with aging^8,11,12,122^, a number of cellular processes, including chronic inflammation, dysregulated proteostasis, synaptic function, retrotransposon activation, and genomic instability, are altered with aging and can impact cognitive function and memory formation^15,107,123–125^. Although there are sex differences in aging-related memory deficits and Alzheimer’s disease^2–6^, this is the first study, to our knowledge, that examines how aging-related changes in alternative splicing and chromatin accessibility in the hippocampus differ between males and females.

Using a genome-wide approach to understand how aging and sex affect the transcriptome and chromatin accessibility in the mouse hippocampus, our results provide a framework of the molecular mechanisms that may contribute to aging-related memory impairment. Our findings support a number of themes that are consistent with existing hypotheses of aging, including aging-related inflammation, changes in myelination, synaptic alterations, calcium dysregulation, loss of heterochromatin and increased retrotransposon activity^15,107,123–128^. Our results also, however, reveal significant sex differences in the transcriptome and chromatin environment in the hippocampus, and further show that aging amplifies sex bias in expression and chromatin accessibility.

### Aging-related changes in immune genes

Immune response genes were upregulated with aging in both male and female hippocampus, in line with numerous reports of chronic inflammation, upregulation of immune genes and activation of microglia with aging in the hippocampus and other brain regions^9,11,12,22,127,129–132^. We found more changes in gene expression overall with aging in female hippocampus than male hippocampus (Fig. 1D), a result that is consistent with previous microarray studies in both human and mouse hippocampus^22,130,131^, but that differs from other brain regions^130^. In agreement with the fact that females mount stronger immune responses than males^133,134^, we found that female hippocampus showed larger aging-related changes in expression of immune-response genes (e.g. *C4b*, Fig. 3D). Given that the hippocampus has been reported to be more immune-alert than other brain regions^135^, the greater aging-related changes in gene expression in female hippocampus may result in part from higher female immune response.

### Aging-related changes in myelin sheath genes

We detected significant aging-dependent changes in the expression and alternative splicing of myelin sheath genes. Produced by oligodendrocytes, myelin is important for axon integrity and action potential conductance^136^, and has been found to be critical for memory retention^137^. The process of remyelination in response to injury involves an inflammatory response mediated by astrocytes and microglia, which leads to myelin debris clearance and OPC recruitment and differentiation into mature oligodendrocytes that in turn, generate new myelin sheath^128,138^. During aging, impaired myelin debris clearance and diminished OPC differentiation reduces the efficiency of remyelination^128,139–141^. A previous study showed that enhancing OPC proliferation and myelination in the aged hippocampus increased memory retention in mice^142^. We detected aging-dependent female bias in the expression of genes encoding myelin sheath components including *Mag*, *Mbp*, *Mog* and *Mal* (Fig. 2D). The compact myelin gene, *Mbp,* was upregulated with aging in female but not male hippocampus by RNA-seq and qPCR, although MBP protein did not show sex-bias in overall abundance (Fig. 4C). There were, however, sex differences in the abundance of specific MBP isoforms in aged hippocampus (Fig. 4C and Supp. Fig. 1D,E), indicating that male and female hippocampus show differential regulation of aging-related *Mbp* transcript and protein isoform expression. Altogether, the female-biased expression of myelin sheath-related genes with aging suggests that female hippocampus undergoes less myelin degeneration or more remyelination than male hippocampus during aging, consistent with studies in aged rats showing a female bias in remyelination and white matter volume^57,58^. Indeed, the female-biased aging-related upregulation of *Bcas1*, a marker of early actively myelinating oligodendrocytes^143^, could reflect more active remyelination in aged female versus male hippocampus. Because an immune response is required for myelin debris clearance and OPC recruitment^128,138^, one possibility is that higher immune response in female hippocampus may lead to more efficient myelin debris clearance and OPC recruitment necessary for remyelination.

We found aging-related splicing differences not only in *Mbp*, but also in myelin sheath genes *Mag* and *Bcas1* in male and female hippocampus (Fig. 4B). The alternative splicing of *Mag,* which encodes a myelin-membrane protein, into large (L-MAG), exon 12-skipped, and small (S-MAG), exon 12-included isoforms is tightly regulated during development, with L-MAG predominating during early myelinogenesis and S-MAG expression increasing during maturation^79,144^. The MAG isoforms associate with different signaling pathways, leading to functional differences; L-MAG is more involved in initial myelination with potentially larger myelination capacity and S-MAG is involved in maintaining myelination^144^. Our finding that aging resulted in an increase in S-MAG and decrease in L-MAG isoforms could point to aging-related reduced myelin formation or capacity. Similarly, the alternative splicing of exon 2 in *Mbp* is also known to be regulated during development^145,146^. MBP isoforms containing exon 2 are enriched during active myelin formation whereas isoforms excluding exon 2 are predominately expressed in adulthood^80,82^. These isoforms differ in their subcellular location and function, with exon 2-containing isoforms localized to the cytosol and nucleus where they regulate myelin development and differentiation, whereas exon 2-excluding isoforms are localized to the oligodendrocyte myelin membrane where they are essential for the formation and function of compact myelin^145^. In rats, loss of exon 2-included isoforms (but not other MBP isoforms) with aging was associated with split and retracted myelin sheaths especially at paranodal regions^83^. Our finding that exon 2-containing MBP isoform expression decreased with aging suggests that this decrease may result in the loss of molecular signaling pathways regulating remyelination and maintenance of the axo-glial junction. Less is known about alternative splicing of the myelination gene, *Bcas1,* however aging-related splicing of *Bcas1* has been previously reported in mouse hippocampus (sex not specified)^11^. Notably, in oligodendrocytes the alternative splicing of *Mag*, *Mbp*, and *Bcas1* is regulated by the RNA binding protein quaking^84,147,148^. Although we did not detect aging-related differential expression or alternative splicing of the quaking gene (*Qk/Qki*), phosphorylation of quaking can regulate its binding to and stabilization of *Mbp* RNA^149^. Altogether, it is possible that altered quaking function during aging in the hippocampus may lead to aging-related changes in alternative splicing of *Mag*, *Mbp* and *Bcas1*, contributing to the decrease in myelination capacity observed with aging.

### Aging-related changes in synaptic function genes

Previous work has shown that in the hippocampus, aging is associated with alternations in synaptic function, including deficits in synaptic plasticity induction, reduction in postsynaptic density size and dysregulated calcium currents^14,15,150^. In alignment with these studies, we detected changes in the expression and alternative splicing of genes regulating synaptic function with aging in the hippocampus of male and female mice (Tables 1 and 2). We found aging-related changes in the expression of genes encoding both inhibitory and excitatory synaptic-related proteins, as well as changes in calcium buffer and calcium channel genes. We found several L- and T-type calcium channel subunit genes that were either downregulated with aging or showed aging-related changes in alternative splicing. Calcium is required for long-lasting synaptic plasticity in the hippocampus, and is tightly controlled in neurons through regulation of calcium buffers, calcium release from internal stores, as well as calcium influx through NMDA receptors and voltage-dependent calcium channels^126^. Dysregulated calcium homeostasis not only suppresses synaptic plasticity, but also causes neurons to be more vulnerable to damage from oxidative stress, and previous work has shown in the hippocampus of rodents, aging results in increased L-type voltage-dependent calcium currents that disrupt long-lasting synaptic plasticity and memory^121,126,150,151^. Additional genes important for synaptic plasticity and memory, including *Camk2a*^152^, *Ablim3*^153^ and *Cplx2* were downregulated with aging in both sexes. Furthermore, male and female hippocampus showed different aging-related changes in expression and alternative splicing, including male-biased expression of the cysteine/glutamate antiporter gene *Slc7a11/xCT*, whose deletion in male mice has been shown to improve aging-related memory deficits^154^.

### Sex differences in gene expression and splicing

Sex-biased expression and splicing was observed in both young and aged hippocampus. Sex differences in the brain can result from the influence of gonadal sex hormones and sex chromosome-expressed genes. In both humans and rodents, circulating and brain-derived estrogens as well as circulating androgens diminish with aging, and their loss is linked to memory impairment^155–159^. We found that relatively few genes showed sex-biased expression in young adult hippocampus (Fig. 2A), and most were sex chromosome-expressed genes. Two notable exceptions were the male-biased expression of synaptic vesicle gene, complexin 2 (*Cplx2*), and the female-biased expression of the neuropeptide gene, *Npy. Npy* has been shown to be upregulated by estradiol in the hippocampus^160^, and correspondingly, we found female-biased expression in young adult hippocampus but aging-related downregulation in female hippocampus. Sex chromosome-expressed genes themselves can also contribute to downstream sex-biased expression. For example, the X chromosome-expressed genes *Kdm6a/Utx* and *Kdm5c* are histone demethylases that show female-biased expression, regulate transcription in the brain, and show incomplete functional overlap with their Y-chromosome paralogs *Uty* and *Kdm5d*^161–164^. Therefore, sex hormones and sex chromosome expressed genes both likely influence the sex-biased expression of hippocampal genes we observe.

Sex-biased alternative splicing can also lead to downstream sex-bias in the transcriptome of the hippocampus. We found sex-biased alternative splicing of *Hmga1* with male-biased inclusion of the splice isoform corresponding to long exon-containing *Hmga1a* over the short exon-containing *Hmga1b* (Fig. 5B). HMGA1 is a chromatin modifier and splicing protein whose binding results in more open chromatin^165,166^. While the function of the different splice isoforms in chromatin accessibility is not understood, previous work in female human cell lines has shown that *Hmga1a*, but not *Hmga1b*, regulates the splicing pattern of presenilin-2 seen in sporadic Alzheimer’s disease^167^. HMGA1a has also been shown to regulate the splicing and DNA-binding of estrogen receptor α in breast cancer cells^168,169^. Another gene known to regulate splicing, *Miat/Gomafu*, also exhibited sex-biased alternative splicing in young adult hippocampus (Fig. 5B). *Gomafu* expression has been shown to be regulated by estrogen in breast cancer cells^170^ and by synaptic activity in neurons^86,87^, and although many alternatively spliced *Gomafu* isoforms have been identified, little is known about their distinct functions^86^. Therefore, the interplay of sex hormones and sex-biased expression and alternative splicing of regulatory factors may contribute to downstream sex-biased patterns in the hippocampus.

### Aging-related changes in chromatin accessibility

This is the first study, to our knowledge, that examines aging-related changes in chromatin accessibility in the hippocampus of both males and females. Our finding that chromatin structure is more open with aging in both sexes (Fig. 6C,D) is consistent with previous findings in male hippocampal neurons^16^ and with the “heterochromatin loss” model of aging^125,171,172^. This model suggests that there is a loss of heterochromatin during aging, leading to alterations in the expression of genes residing in these regions^172^. Chromatin accessibility is associated with active regulatory elements and is regulated by chromatin and histone modifiers. Histone variants and histone post-translational modifications are associated with open or closed chromatin, and in general, CpG island methylation is correlated with decreased chromatin accessibility^113,114,173^. Changes in the expression of histone proteins, as well as altered histone modifications and DNA hypo- or hyper-methylation are widely documented to occur during aging; indeed, aging-related changes in DNA methylation serve as the basis of epigenetic clocks^124,125,174,175^. Therefore, loss of histone proteins, loss of repressive histone marks and changes in DNA methylation likely contribute to the aging-related opening in chromatin accessibility we detected in the male and female hippocampus.

Our results show that most of the aging-related increases in chromatin accessibility occurred in intergenic and intronic regions, many of which contained retrotransposon LINE-1-derived sequences (Fig. 7C). Furthermore, we found increased expression of LINE-1 transcripts with aging in the hippocampus (Fig. 7E). Increased retrotransposon activity has been reported to accompany aging in both humans and rodents, and the brain has been reported to exhibit more retrotransposon activity than other tissues^16,18,107,176^. In young mammalian cells, the histone deacetylase SIRT6 binds to LINE-1 retrotransposon sequences and represses their activity by modulating the heterochromatin status of nearby regions. However, during aging, SIRT6 is depleted from LINE-1 regions, and histone repressive marks and CpG methylation is lost in transposable element regions, leading to expression of LINE-1 transcripts and the accompanying DNA damage and cellular immune activation. Furthermore, repression of aging-related retrotransposon activity through overexpression of SIRT6 have been shown to increase lifespan^105,107,125^. We found that full-length intact LINE-1 regions in both male and female hippocampus remained robustly inaccessible with aging (Supp. Fig. 3C), suggesting that transposition-capable sequences as a whole remain suppressed, although individual instances of full-length LINE-1 element activity may occur. LINE-1 derived sequences that opened with aging were truncated sequences incapable of retrotransposition, however expression of truncated LINE-1 transcripts have been reported to cause cellular inflammation and DNA damage^102–104^. Thus, loss of repressive histone marks and CpG methylation at LINE-1 regions likely contributed to the aging-related opening of chromatin at those regions and to the increased LINE-1 expression we observed in male and female hippocampus.

### Sex differences in chromatin accessibility

We found that male and female hippocampus showed sex differences in chromatin accessibility on autosomes, and the number of regions with sex-biased accessibility increased with aging (Fig. 8D). These differences could be a result of sex-biased expression or activity of histone and/or DNA methylation modifiers. Sex hormones have been shown to affect DNA methylation in human blood^177^ and chromatin accessibility in female mouse young adult ventral hippocampus^178^. We detected few sex differences in chromatin accessibility in young adult hippocampus (Fig. 8A); however, differences in aging-related DNA methylation between sexes could contribute to the sex-biased accessibility observed in aged hippocampus. Indeed, it has been reported that aging-related DNA methylation changes differed between male and female hippocampus in mice^179^. Furthermore, studies in human blood cells have shown aging-related sex differences in DNA methylation^180–182^. We found male-biased chromatin accessibility in a subset of promoters and CpG-rich regions (Fig. 8C,D; Supp. Fig. 5); raising the possibility that CpGs in these regions could be hypermethylated in female compared to male hippocampus. We did not detect sex-bias in the expression of DNA methylation-related proteins, therefore DNA methylation patterns established in development, or other mechanisms regulating the activity of DNA methylation-related proteins, may contribute to these differences in accessibility.

Alternatively, or in conjunction with sex-biased DNA methylation, sex differences in histone modifications may contribute to the male-bias accessibility in promoter regions we observed. As mentioned previously, we detected female-biased expression of the X chromosome-linked histone modifiers *Kdm6a* and *Kdm5c*. *Kdm5c* is a histone demethylase that removes histones from H3K4me2/3^161,183,184^. H3K4me3 is a histone mark associated with active promoters, and *Kdm5c* has been found to reduce H3K4 trimethylation at promoters and suppress transcription^161,162,185^. *Kdm5c* escapes X-inactivation and has been shown to exhibit female-biased expression in the brain^186^. We found that *Kdm5c* showed female-biased expression in both young adult and aged hippocampus. In young adult and aged males, the Y-chromosome paralog, *Kdm5d*, was expressed at slightly lower levels than *Kdm5c*. Although the summed *Kdm5c/d* expression in males is higher than *Kdm5c* expression in female hippocampus, *Kdm5d* is unable to compensate for *Kdm5c* function in the brain^187^. Indeed, loss-of-function mutations in *Kdm5c* reduce H3K4me2/3 demethylation in a subset of promoters and enhancers, and have been shown to lead to a form of X-linked intellectual disability in males^183,188^. *Kdm5c* shows preferential binding to promoters, especially those containing CpG islands^162^, and we found male-biased accessibility at promoters and CpG-rich regions. Therefore, one possibility is that female-biased expression of *Kdm5c* may result in greater demethylation of H3K4me3 at promoters in female hippocampus and contribute to lower chromatin accessibility at those regions. A previous study of male patients with X-linked intellectual disability and mutations in *Kdm5c* found promoter CpG sites with reduced DNA methylation in the blood of patients compared to controls^189^. Furthermore, the authors showed that these promoter regions also showed female-biased DNA hypermethylation in the brain of controls, as well as DNA methylation that correlated with *Kdm5c*/*Kdm5d* dosage in individuals with karyotypes of varying number of X chromosomes^189^. Therefore, female-biased *Kdm5c* expression may lead to greater histone demethylation and DNA methylation at promoters in females, thereby contributing to the male-bias in promoter accessibility we observed. Notably, the male-bias in promoter accessibility is not sufficient to drive sex-biased expression of the associated genes. Therefore, other mechanisms may compensate for sex differences in promoter accessibility to regulate expression.

In conclusion, this study provides a comprehensive map of gene expression, splicing and chromatin accessibility changes in the hippocampus during aging in both male and female mice. Our findings of sex-biased alternative splicing and aging-dependent sex-bias in expression and chromatin accessibility illustrate the importance of including female animals in studies of brain aging. These findings also provide novel starting points for future research of chromatin structure and transcriptomics in the hippocampus that can guide treatments for aging-related memory decline.

## Acknowledgement

The authors would like to thank UCLA Broad Stem Cell Research Center Sequencing Core and UCLA Neuroscience Genomic Core for data collection and Yue Qin for data analysis support. This research was supported by funding from the David Geffen Foundation. YT was support by the CSC-UCLA Scholarship by China Scholarship Council, Whitcome Pre-doctoral Fellowship and Dissertation Year Fellowship from UCLA.

---

**Supplemental Figure 1:**
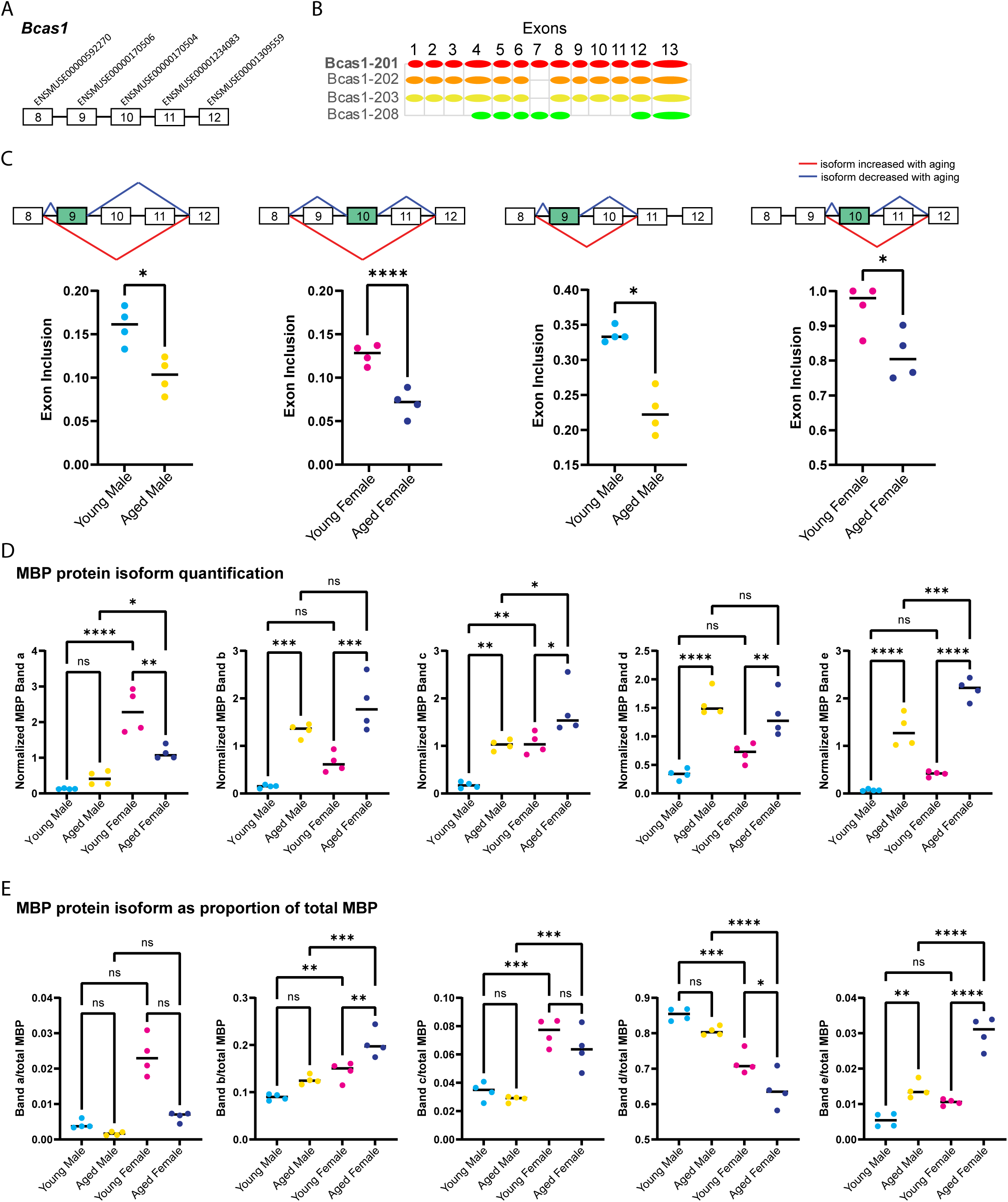
Aging-associated alternative splicing of myelin sheath genes *Bcas1* and *Mbp*. (A) *Bcas1* exons involved in aging-related alternative splicing events in the hippocampus. (B) *Bcas1* protein-coding splice variants from Ensembl^190^ release 102 with a diagram of included exons. Bcas1-201 is bold to indicate Ensembl’s designation as a stable, reviewed, and high quality transcript annotation. (C) Diagram and exon inclusion values for each *Bcas1* aging-related alternative splicing event determined by rMATS. In the first two panels, skipping of exons 9-11 is depicted (copy of Fig. 4A *Bcas1* panels). For male and female hippocampus, one additional aging-related alternative splicing event was detected for each (right two panels), in which exons 9 and 10 were skipped with aging. The alternative splicing in all these events skip exon 9 or exon 10, and it is unclear if these events defined by rMATS represent the increasing expressing of isoform Bcas1-208 versus Bcas1-201 with aging in both males and females, or if other unannotated *Bcas1* transcripts are produced. * indicates p < 0.05, *** indicates p < 0.001. (D) Western blot analysis for MBP protein isoforms in young adult and aged hippocampus from Fig. 4C. MBP band “a” young female vs aged female p = 0.001; young male vs young female p < 0.0001; aged male vs aged female p = 0.043. MBP band “b” young male vs aged male p < 0.001; young female vs aged female p < 0.001. MBP band “c” young male vs aged male p = 0.008; young female vs aged female p = 0.027; young male vs young female p = 0.006; aged male vs aged female p = 0.020. MBP band “d” young male vs aged male p < 0.0001; young female vs aged female p = 0.010. MBP band “e” young male vs aged male p < 0.0001; young female vs aged female p < 0.0001; aged male vs aged female p < 0.001. ANOVA with Sidak’s multiple comparison test was used for all tests. * indicates p < 0.05, ** indicates p < 0.01, *** indicates p < 0.001, **** indicates p < 0.0001 and ns indicates not significant. (E) Same as in D, except each band was normalized to total MBP to determine how each isoform’s relative proportion changed with aging. MBP band “a” p > 0.05. MBP band “b” young female vs aged female p = 0.003; young male vs young female p = 0.006; aged male vs aged female p < 0.001. MBP band “c” young male vs young female p < 0.001; aged male vs aged female p < 0.001. MBP band “d” young female vs aged female p = 0.023; young male vs young female p < 0.001; aged male vs aged female p < 0.0001. MBP band “e” young male vs aged male p = 0.003; young female vs aged female p < 0.0001; aged male vs aged female p < 0.0001. ANOVA with Sidak’s multiple comparison test was used for all tests.

**Supplemental Figure 2:**
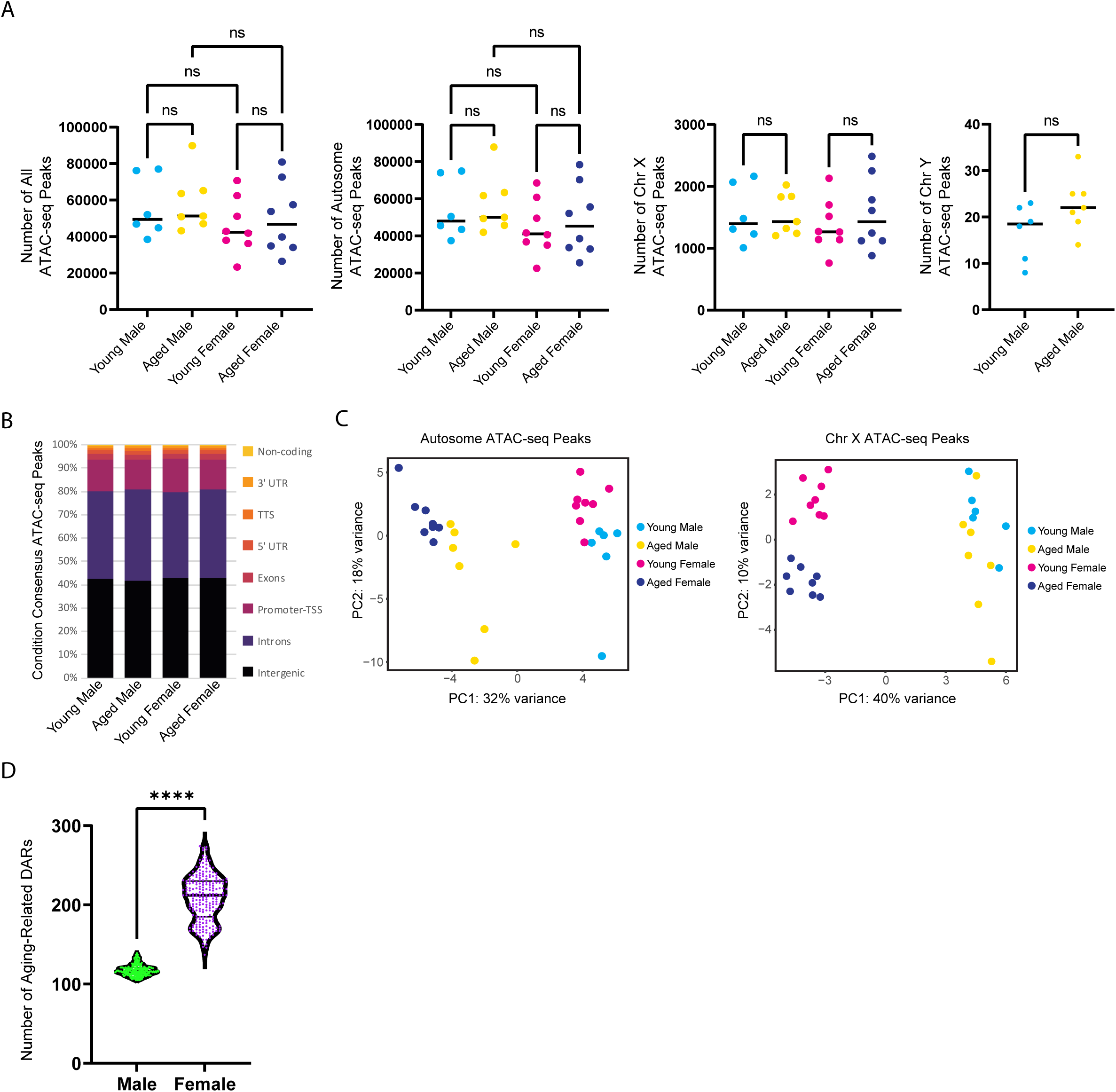
Number and annotation of ATAC-seq peaks are similar across conditions. (A) From left to right, number of ATAC-seq peaks with FDR < 0.05 located on all chromosomes, autosomes, chromosome X or chromosome Y in the hippocampus of 6 young adult male, 7 aged male, 8 young adult female and 8 aged female mice. All peaks: Kruskal-Wallis with Dunn’s multiple comparisons test adjusted p > 0.05 for all comparisons. Autosome peaks: one-way ANOVA with Sidak’s multiple comparison test adjusted p > 0.05 for all comparisons. Chromosome X peaks: one-way ANOVA with Sidak’s multiple comparison test adjusted p > 0.05 for both comparisons. Chromosome Y peaks: unpaired t-test p > 0.05. ns = not significant. (B) Genome annotation of condition consensus ATAC-seq peaks. These peaks were generated from sample peaks sets and merged to generate the total consensus peak set. (C) Principle component analysis of ATAC-seq fragments in the total consensus peaks located on either autosomes (left) or chromosome X (right). There were too few total consensus ATAC-seq peaks located on chromosome Y to perform principal components analysis. (D) Number of significant aging-related differentially accessible regions (DARs) in male and female hippocampus for 224 combinations of samples which included all 6 young male and all 7 aged male samples with different combinations of 6 young female and 7 aged female samples; Mann-Whitney U = 2, **** indicates p < 0.0001.

**Supplemental Figure 3:**
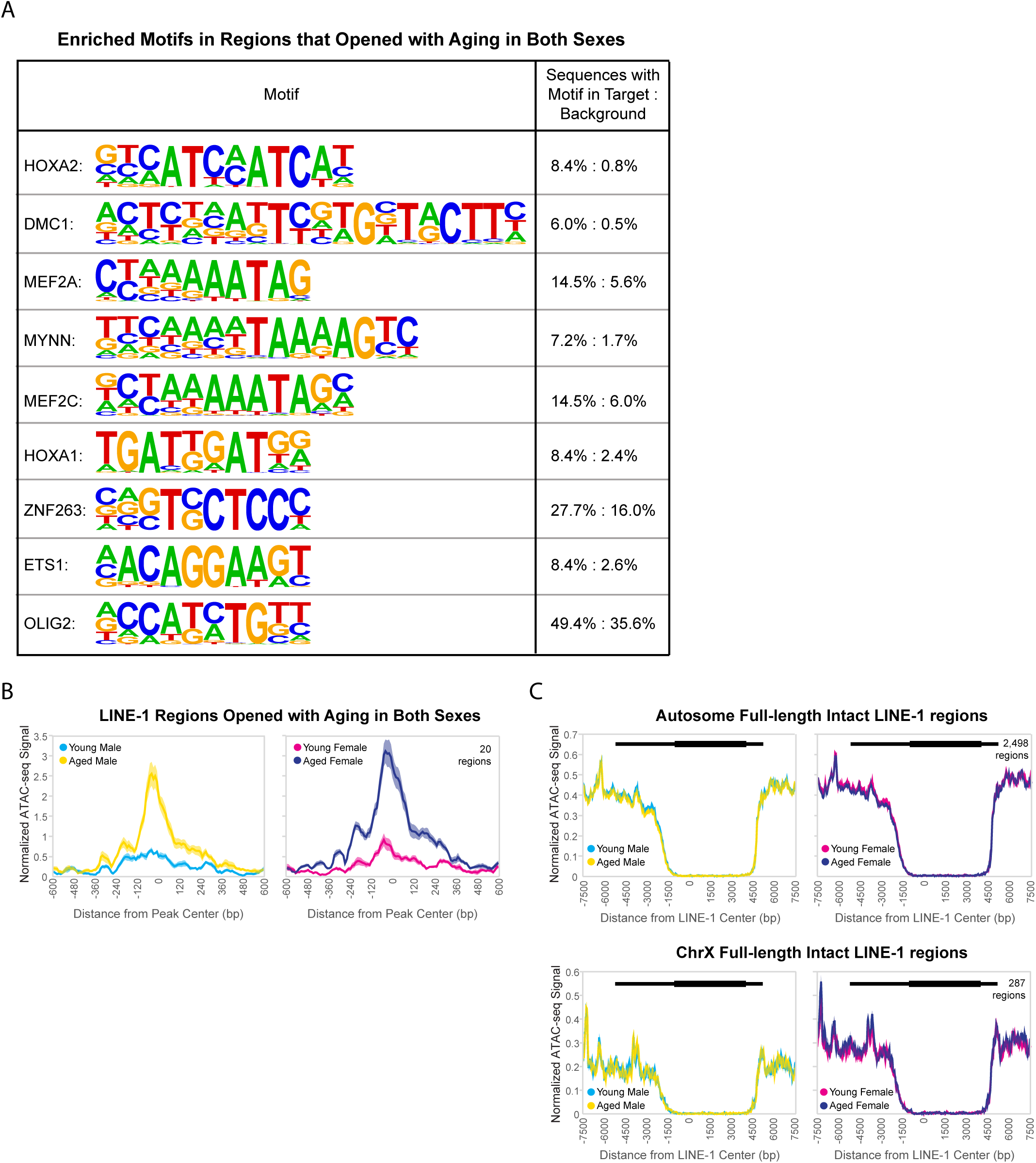
Aging-associated changes in chromatin accessibility. (A) Motifs enriched in regions that gained accessibility with aging in both female and male hippocampus (FDR < 0.05). (B) ATAC-seq profiles for regions containing LINE-1 elements that showed increased accessibility with aging in both males and females (FDR < 0.05). Solid lines indicate the average of each condition’s normalized histogram (n = 6 young adult male, 7 aged male, 8 young adult female, 8 aged female) with shading indicating s.e.m. (C) ATAC-seq profiles (as in B) for full-length intact LINE-1 elements on autosomes (above) and on the X chromosome (below). Shown above each ATAC-seq plot is an aligned schematic of a full-length LINE-1 element.

**Supplemental Figure 4:**
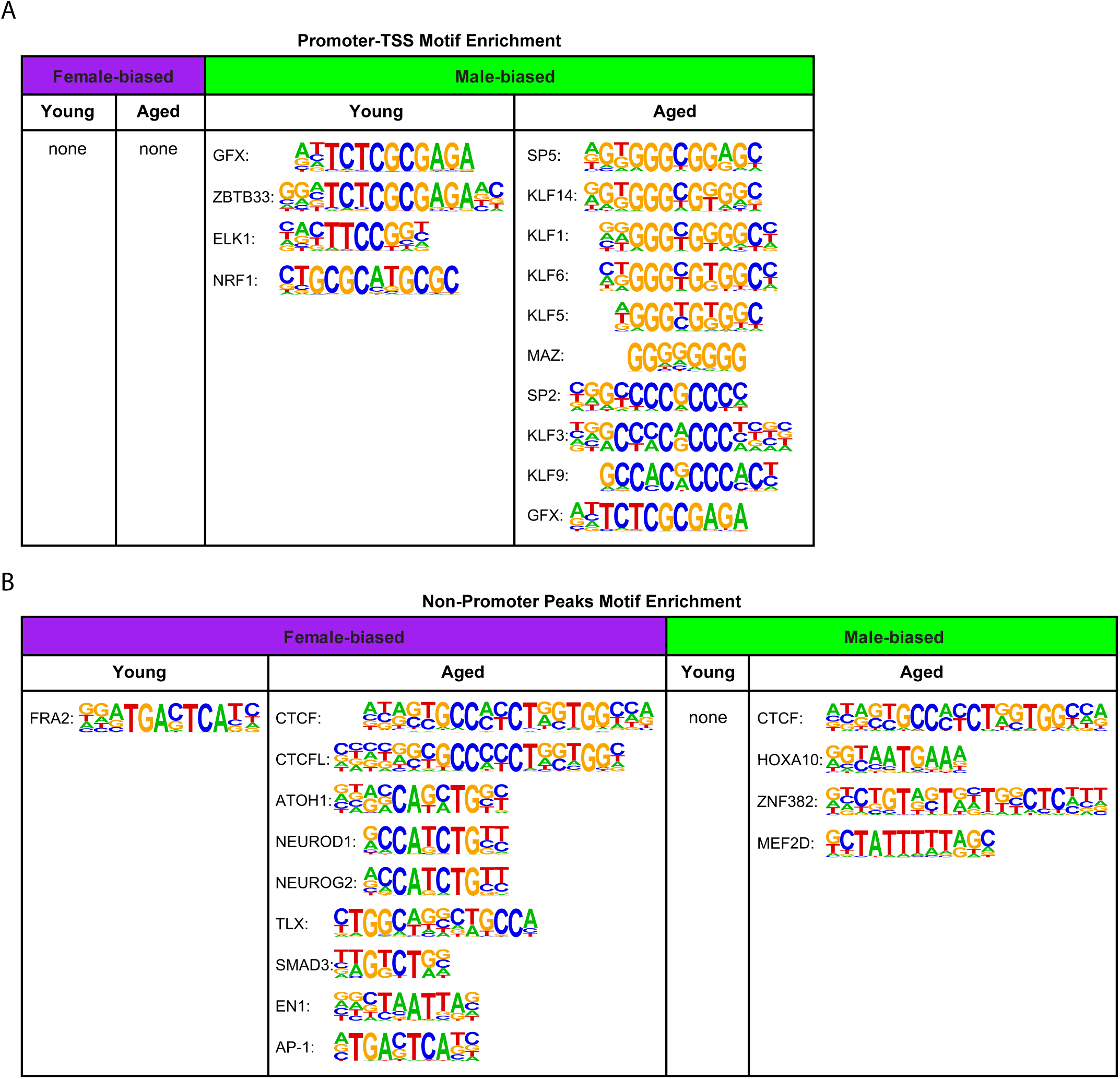
Motifs in regions with sex-bias in chromatin accessibility. (A) Names and motifs enriched in differentially accessible autosome promoter-TSS regions between female and male hippocampus (FDR < 0.05). Zero and three autosome promoter-TSS regions showed a female-bias in accessibility in young adult and aged hippocampus, respectively, yielding no significantly enriched motifs. There were 26 motifs significantly enriched in regions showing male-bias in aged hippocampus, and the top 10 most significant are shown. (B) Name and enriched motifs as in (A), but for non-promoter regions that were differentially accessible between female and male hippocampus. No enriched motifs were found for non-promoter regions that showed a male-bias in accessibility in young animals.

**Supplemental Figure 5:**
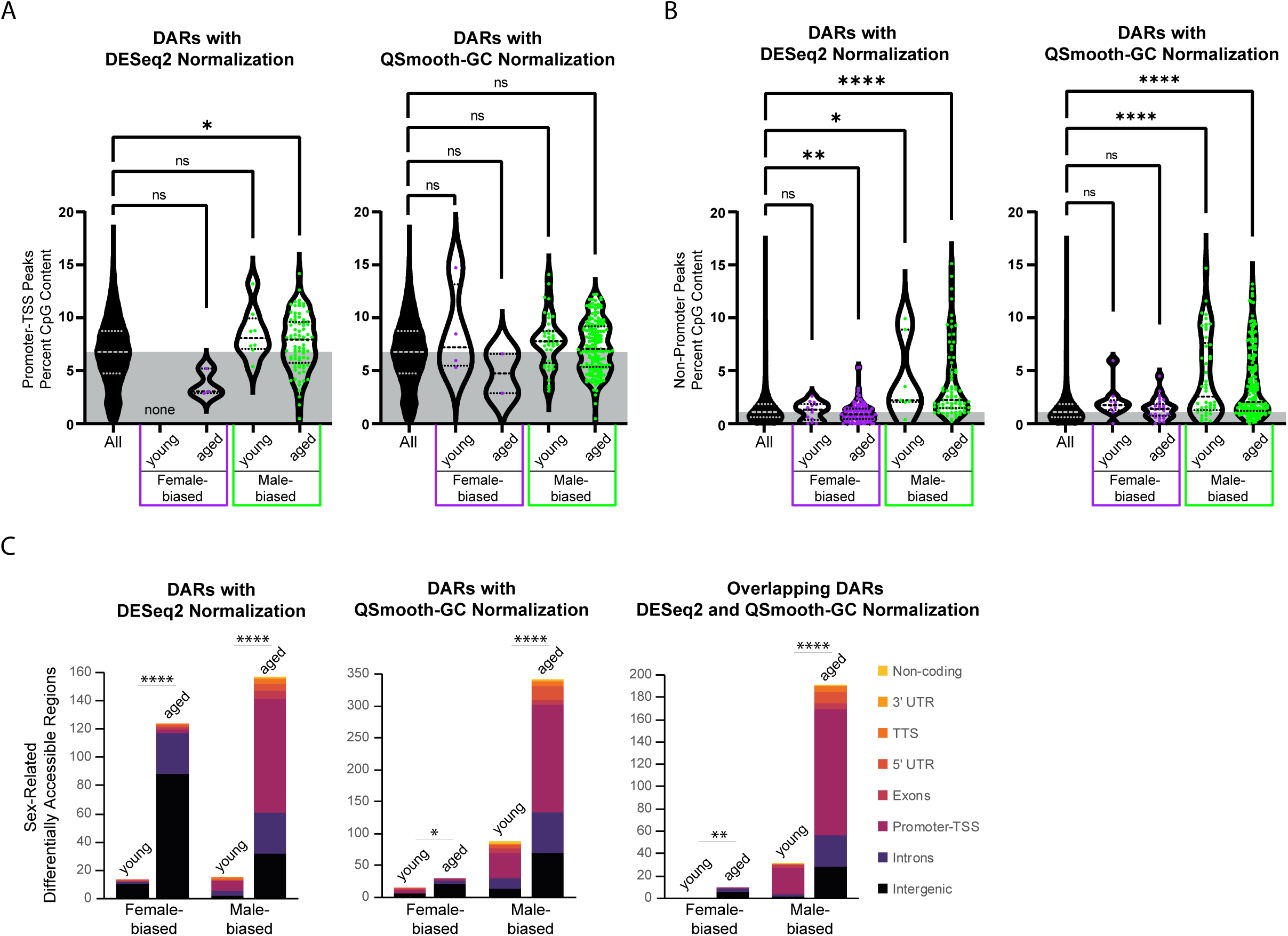
Male bias in chromatin accessibility at promoters and CpG-rich regions. (A) Violin plots of ATAC-seq autosome promoter-TSS regions’ CpG content. CpG content was calculated using Homer^39^ for all ATAC-seq promoter-TSS regions (All; 9,541 regions) and compared to those autosome promoter regions that were differentially accessible with DESeq2 normalization (left panel) or QSmooth-GC normalization (right panel). For data in both panels, Kruskal-Wallis tests with Dunn’s multiple comparison tests were used. Average ± s.e.m., All promoter ATAC-seq regions CpG = 6.7% ± 0.0. For autosome promoter DARs with DESeq2 normalization: more open in young adult female (0), more open in aged females (3), more open in young adult males (8) and more open in aged males (80); open aged female CpG = 3.7% ± 0.8, p = 0.18; open young male CpG = 8.5% ± 0.8, p = 0.27; open aged male CpG = 7.6% ± 0.3, p = 0.02. For autosome promoter DARs with QSmooth-GC normalization: more open in young females (4), more open in aged females (2), more open in young males (41) and more open in aged males (167); more open young female CpG = 8.6% ± 2.1, p = 1; more open aged female CpG = 4.7% ± 1.9, p = 1; more open young male CpG = 7.8% ± 0.4, p = 0.10; more open aged male CpG = 7.3% ± 0.2, p = 0.09.). (B) CpG content as in (A), but for non-promoter regions, all ATAC-seq non-promoter regions (All; 82,692 regions) CpG = 1.6% ± 0.0 (average ± s.e.m.). For data in both panels, Kruskal-Wallis tests with Dunn’s multiple comparison tests were used. For non-promoter autosome DARs with DESeq2 normalization, more open in young females (14), more open in aged females (121), more open in young males (7) and more open in aged males (77); open young female CpG = 1.2% ± 0.2, p = 1; open aged female CpG = 1.0% ± 0.1, p = 0.002 ; open young male CpG = 4.2% ± 1.4, p = 0.04; open aged male CpG = 3.8% ± 0.4, p < 0.0001. For non-promoter DARs with QSmooth-GC normalization: more open in young females (10), more open in aged females (27), more open in young males (46) and more open in aged males (174); more open young female CpG = 2.0% ± 0.5, p = 0.28; more open aged female CpG = 1.4% ± 0.2, p = 1; more open young male CpG = 4.4% ± 0.6, p < 0.0001; more open aged male CpG = 3.7% ± 0.2, p < 0.0001. (C) Genome annotation of autosome differently accessible regions (DARs) between female and male hippocampus. Left panel shows annotation of regions found to be differentially accessible using DESeq2 normalization (copy of Fig. 8D). Chi-square test, young vs aged *X^2^*= 87.7 more open female, *X^2^* = 117.3 more open male, p > 0.00001. Middle panel shows annotation of regions found to be differentially accessible using QSmooth-GC Normalization (FDR < 0.05). Chi-square test, young vs aged *X^2^* = 5.2 more open female, p = 0.021; *X^2^* = 151.1 more open male, p < 0.0001. Right panel shows annotation of regions found to be differentially accessible using either DESeq2 or QSmooth-GC Normalization (FDR < 0.05). Chi-square test, young vs aged *X^2^* = 7.4 more open female, p > 0.01, *X^2^* = 113.5 more open male, p < 0.00001. * indicates p < 0.05, ** indicates p < 0.01, **** indicates p < 0.0001.

**Supplemental Figure 6:**
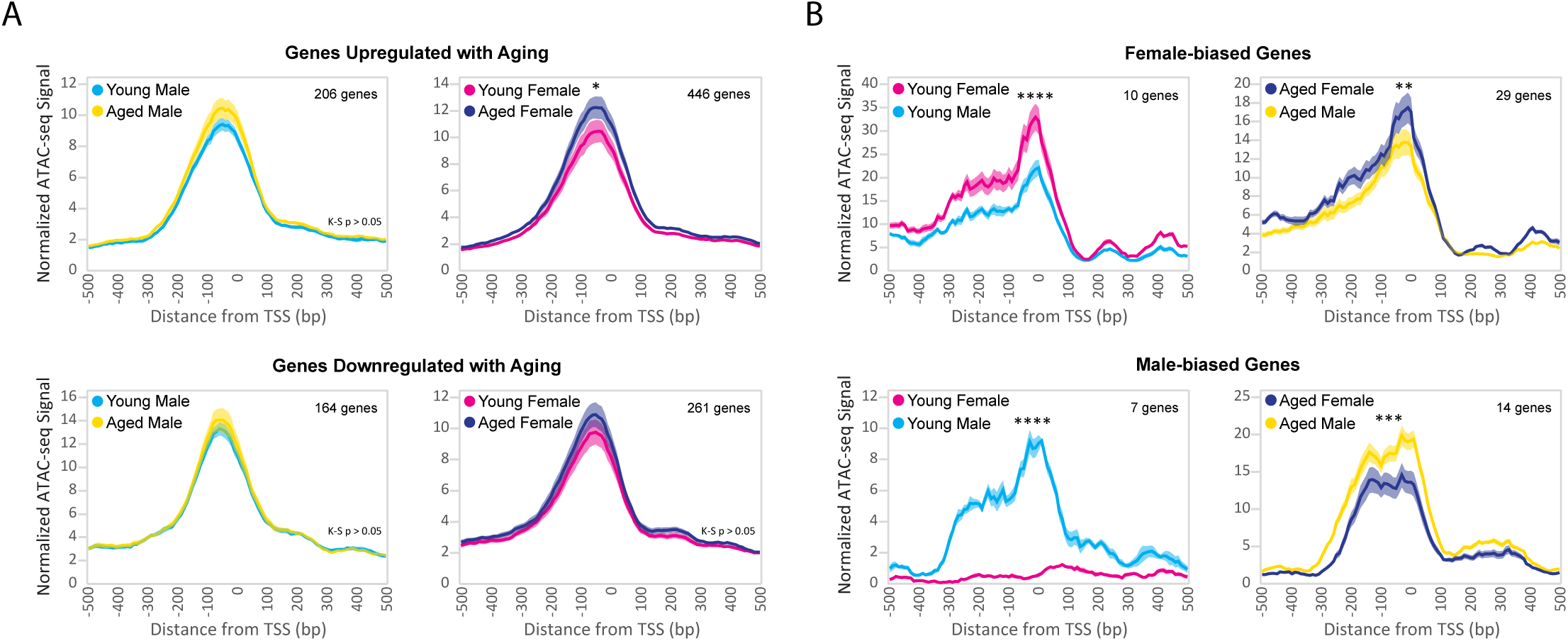
Aging and sex-related differential expression correlates with ATAC-seq signal at gene TSSs. (A) ATAC-seq profiles surrounding the TSS of genes that were upregulated with aging (top row) or downregulated with aging (bottom row). Solid lines indicate the average of each condition’s normalized histogram (n = 6 young adult male, 7 aged male, 8 young adult female, 8 aged female) with shading indicating s.e.m. (upregulated with aging in male: Kolmogorov-Smirnov D = 0.17, p = 0.11; female: Kolmogorov-Smirnov D = 0.21, p = 0.03; downregulated with aging in male: Kolmogorov-Smirnov D = 0.11, p = 0.59; female: Kolmogorov-Smirnov D = 0.18, p = 0.08). (B) ATAC-seq profiles surrounding the TSS of genes that were higher expressed in female hippocampus (top row) or higher expressed in male hippocampus (bottom row). Solid lines indicate the average of each condition’s normalized histogram (n = 6 young male, 7 aged male, 8 young female, 8 aged female) with shading indicating s.e.m. (higher expressed in female young: Kolmogorov-Smirnov D = 0.26, p = 0.003; aged: Kolmogorov-Smirnov D = 0.22, p < 0.0001; higher expressed in male young: Kolmogorov-Smirnov D = 0.82, p < 0.0001; aged: Kolmogorov-Smirnov D = 0.31, p = 0.0001).* indicates p < 0.05, ** indicates p < 0.01, *** indicates p < 0.001, **** indicates p < 0.0001.

**Supplemental Table 1:** RNA-seq DE data. RNA-seq gene data from young adult and aged, female and male hippocampus.

**Supplemental Table 2:** Alternative splicing data. Alternative splicing events from young adult and aged, female and male hippocampus. SE = skipped exon, A5SS = alternative 5’ splice site, A3SS = alternative 3’splice site, MXE = mutually exclusive exons, RI = retained intron.

**Supplemental Table 3:** ATAC-seq data. ATAC-seq total consensus peaks data from young adult and aged, female and male hippocampus.

